# Intrinsic regulation of intestinal stem cell fate and homeostasis by Tet

**DOI:** 10.64898/2026.05.05.722984

**Authors:** Niccole Auld, Ye-Jin Park, Tyler Jackson, Sarah Eleraky, Yara Younes, Chung-Yi Liang, Tzu-Chiao Lu, Shangyu Gong, Zhiyong Yin, Bo Sun, Yulong Zhang, Tao P. Wu, Yanyan Qi, Hongjie Li

## Abstract

How developmental progenitors navigate divergent trajectories to either establish adult stem cell pools or undergo terminal differentiation remains a fundamental gap in stem cell biology. Here, we identify the well conserved Tet protein as an essential transcriptional regulator of intestinal stem cell (ISC) establishment. Developmental Tet depletion causes region-specific ISC loss and compromises adult lifespan, while adult-specific loss drives progressive stem cell exhaustion. Overexpression of Tet leads to ISC expansion in both developing and adult guts. Utilizing a comprehensive single-nucleus transcriptomic atlas spanning gut development, we demonstrate that Tet stabilizes progenitor identity by maintaining epithelial integrity, niche signaling, and fate maintenance. By defining this developmental trajectory, we reveal Tet as a critical factor that drives proper ISC maturation and maintains long-term adult epithelial homeostasis.

## INTRODUCTION

Stem cells must maintain a stable identity while responding to external challenges to sustain epithelial integrity. Disruption of stem cell establishment during development can compromise tissue homeostasis and lead to epithelial dysfunction, including tissue degeneration due to stem cell depletion and cancer driven by aberrant stem cell self-renewal (Barker et al., 2009; Clevers, 2013). The *Drosophila melanogaster* midgut serves as an ideal model system to study these dynamics, as it contains intestinal stem cells (ISCs) and differentiated cell types that are functionally and molecularly analogous to those found in mammals (Micchelli & Perrimon, 2006; Ohlstein & Spradling, 2006; Li & Jasper, 2016; Miguel-Aliaga et al., 2018). Recent work demonstrates that this midgut lineage retains a broadly permissive transcriptional landscape, making cell fates highly plastic and susceptible to conversion if key regulators are lost (X. Guo et al., 2024). This inherent flexibility highlights a critical developmental vulnerability, as progenitors must lock in their identity to prevent premature differentiation.

During embryonic gut development, epithelial lineages arise from a shared endoblast population to produce transient larval cells alongside long-lived adult midgut progenitors (AMPs), which will ultimately expand to establish the mature adult epithelium (Tepass & Hartenstein, 1995; Micchelli et al., 2011; Plygawko et al., 2025). Throughout larval stages, AMPs are maintained in an undifferentiated state within clusters called AMP islands through interactions with Dpp-secreting niche cells called peripheral cells (Mathur et al., 2010). At the onset of metamorphosis, peripheral cells retract their cellular processes and aggregate with other larval cells to form a dense internal structure known as the yellow body (meconium) (Takashima et al., 2011). Meanwhile, the AMPs disperse and migrate outward to establish the nascent adult epithelial layer. Despite this well-defined morphogenetic progression, it remains unclear when and how individual AMPs bifurcate to balance terminal differentiation with establishment of the adult ISC pool.

While many extrinsic signaling pathways regulating adult stem cell behavior have been well defined, including Notch (Micchelli & Perrimon, 2006; Ohlstein & Spradling, 2007; Z. Guo & Ohlstein, 2015; Takashima et al., 2016; Martin et al., 2018), Wnt (Tian et al., 2016; Hu et al., 2021; Y. Wu et al., 2025; Lv et al., 2025), EGFR (Jiang & Edgar, 2009; Biteau & Jasper, 2011; Ngo et al., 2020; Resnik-Docampo et al., 2021), and Hippo (Ren et al., 2010; Poernbacher et al., 2012; Hao et al., 2020), the intrinsic mechanisms that stabilize stem cell fate during development remain poorly understood. To identify regulators of this mechanism, we analyzed the single-cell transcriptomic datasets from recent Fly Cell Atlas studies (Li et al., 2022; Lu et al., 2023; Park et al., 2025) and observed strong enrichment of the *Ten-Eleven Translocation* (*TET*) ortholog, *Tet*, in ISCs and their undifferentiated daughter cells called enteroblasts (EBs) (Fig. S1). While Tet proteins are evolutionarily conserved dioxygenases best known for DNA demethylation (Tahiliani et al., 2009; X. Wu & Zhang, 2017), recent work demonstrates that they also function in an enzyme-independent manner to regulate cell fate decisions (Tsiouplis et al., 2021; Chrysanthou et al., 2022; Stolz et al., 2022). Although *Tet* is a top enriched gene in *Drosophila* ISCs and EBs (Fig. S1), its function has not been studied for fly ISC development or maintenance.

Here, we identify Tet as a critical intrinsic regulator of ISC identity, with stage-dependent requirements. Loss of *Tet* during development results in anterior-specific ISC depletion, increased susceptibility to oxidative stress, and a dramatic reduction in fly lifespan, demonstrating the lasting physiological consequences of impaired stem cell establishment. In adults, *Tet* depletion produces a progressive decline in stem cell maintenance. To define downstream mechanisms, we performed single-nucleus RNA sequencing across five stages spanning midgut development, establishing a comprehensive developing gut atlas (https://hongjielilab.org/gut-atlas/). Our analysis reveals that Tet stabilizes progenitor identity by maintaining a transcriptional network governing epithelial integrity, niche signaling, and fate maintenance. Together, these findings establish Tet as an essential transcriptional regulator that drives proper ISC maturation and maintains long-term adult homeostasis.

## RESULTS

### Tet is required for ISC establishment during development

To test whether Tet is required for stem cell establishment in the gut, we depleted *Tet* using the *esg-Gal4* driver (marked with *UAS-nGFP*), which is continuously expressed in gut progenitor cells from early development through adulthood. Adult flies with *Tet RNAi* eclosed normally but exhibited a pronounced reduction of progenitors in the anterior midgut, with ISC and EB ratio reduced to approximately 2% compared to 14% in control flies (*Luciferase RNAi*) (Fig. 1A-C). In contrast, the posterior midgut showed no significant difference between RNAi and controls. This regional phenotype was recapitulated by multiple independent RNAi lines, demonstrating that the effect is specific to *Tet* loss. To further validate ISC loss in an endogenous context, we crossed the *Tet-T2A-Gal4* (*Tet-Gal4^T2A^*) driver with *Tet RNAi*. We combined this driver with *UAS-mCherry.nls* to track the targeted cells, which revealed a loss of both the *Tet*+ cell population and Delta+ ISCs (Fig. 1D,E).

**Figure 1.**
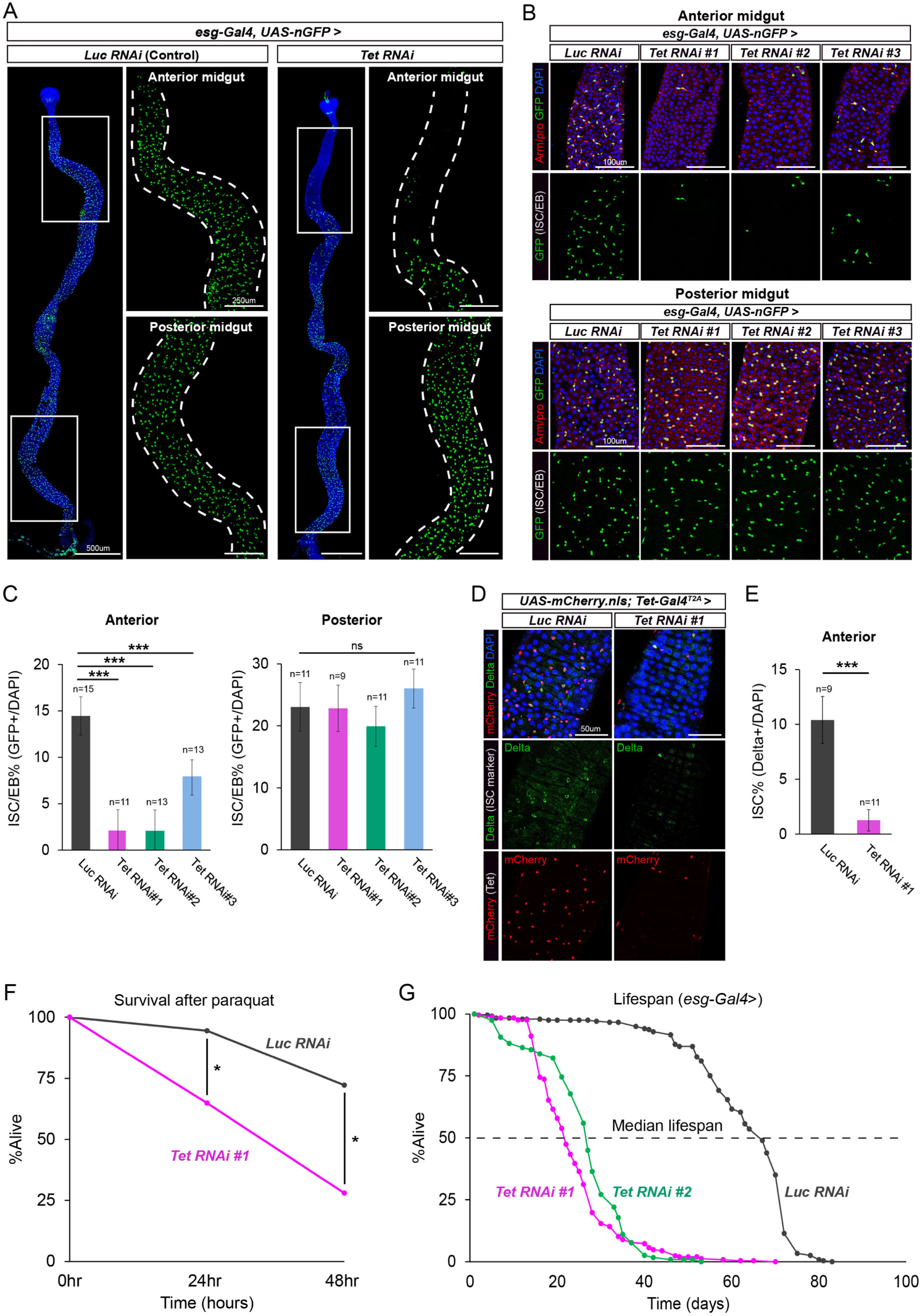
Developmental depletion of *Tet* causes anterior-specific loss of ISCs and reduces adult lifespan. (A) Representative confocal images of whole midguts from control (*Luc RNAi*) and *Tet RNAi* adult flies driven by *esg-Gal4, UAS-nGFP*. Loss of ISCs/EBs (green) is restricted to the anterior midgut. (B) High-magnification images of the anterior and posterior midgut compartments across three independent *Tet RNAi* lines. (C) Quantification of ISC/EB percentage (GFP+/DAPI) in the anterior and posterior midgut. (D) Confirmation of ISC loss using the endogenous *Tet-Gal4^T2A^* driver crossed with *UAS-mCherry.nls* and stained for the ISC marker Delta (green). (E) Quantification of ISC percentage (Delta+/DAPI) in the anterior midgut. (F) Survival curve of adult flies exposed to paraquat-induced oxidative stress over 48 hours. *Tet RNAi* n=54, *Luciferase RNAi* n=53. (G) Lifespan analysis comparing control and *Tet RNAi* adult flies. *Tet RNAi #1* n=247, *Tet RNAi #2* n=118, *Luciferase RNAi* n=240. Lifespans were analyzed with OASIS 2. Bars represent mean ± SD. *p ≤ 0.05, **p ≤ 0.01, ***p ≤ 0.001. p values were from unpaired Student’s t-test.

The regional vulnerability of the anterior midgut aligns with established models of midgut compartmentalization, which demonstrate that anterior and posterior ISCs are transcriptionally and functionally distinct populations governed by differing local signaling environments (Buchon et al., 2013; Li et al., 2013; Marianes & Spradling, 2013). Despite the depletion of anterior ISCs, the overall size and structure of the adult gut remained intact, arguing against widespread cell loss and instead suggesting a shift in cell fate. Given the well-established regional heterogeneity of the midgut, we next focused on mechanistic characterization of anterior ISC loss, while a detailed comparison between anterior and posterior compartments can be addressed in future studies.

### Developmental depletion of *Tet* leads to lasting physiological consequences in the adult

We next examined epithelial organization in day 2 adult midguts following developmental *Tet* depletion. In control flies, anti-armadillo (arm) staining revealed the continuous, honeycomb-like cell junctions characteristic of a healthy epithelium (Resnik-Docampo et al., 2017; Hodge et al., 2023). In contrast, *Tet RNAi* anterior midguts exhibited disrupted junctional continuity and distorted EC morphology, indicative of defective epithelial organization (Fig. S2). Notably, these structural defects were restricted to the anterior midgut, the region where ISC establishment fails. This suggests that the developmental loss of *Tet* not only depletes the stem cell pool but also impairs overall tissue maturation.

Given the combined loss of ISCs and disruption of epithelial architecture, we next asked whether developmental *Tet* depletion impairs adult physiology. Under normal conditions, ISCs proliferate and differentiate in response to stress or age-associated damage to maintain epithelial integrity (Biteau et al., 2008; Amcheslavsky et al., 2009). Consistent with a failure to establish this regenerative capacity, *Tet RNAi* flies exhibited heightened sensitivity to paraquat-induced oxidative stress (Fig. 1F) and a significantly reduced lifespan (Fig. 1G). Together, these findings suggest that impaired ISC establishment and epithelial disorganization caused by *Tet* depletion compromise long-term adult homeostasis and resilience to stress.

### *Tet* expression during development marks the ISC-fated AMPs

To determine when and where Tet functions during gut development, we examined its expression using its endogenous *Tet-Gal4^T2A^* driver across larval, pupal, and adult stages (Fig. 2A). In larval midguts, *Tet* expression was highly enriched in peripheral cells (PCs) but minimally detected within the AMP islands (Fig. 2B). During early metamorphosis the AMP islands break apart and the AMPs migrate outward to form the nascent epithelium while the PCs and larval ECs move inward to form the yellow body (0-18hr after pupa formation (APF)). During this transition, *Tet* expression expanded broadly across the AMP layer (Fig. 2C). This period coincides with extensive progenitor migration, proliferation, and epithelial remodeling, suggesting that *Tet* induction accompanies a highly plastic progenitor state. As development progressed (36-48hr APF), *Tet* expression was selectively retained in a subset of AMPs and downregulated in others (Fig. 2C). From 72hr APF onward, *Tet* expression became restricted to adult ISCs (Fig. 2D-F). Notably, our multi-stage tracking data (summarized in Fig. 2G) indicates that maintaining high *Tet* expression during the mid-pupal stage commits a subset of the progenitor pool to the adult ISC lineage.

**Figure 2.**
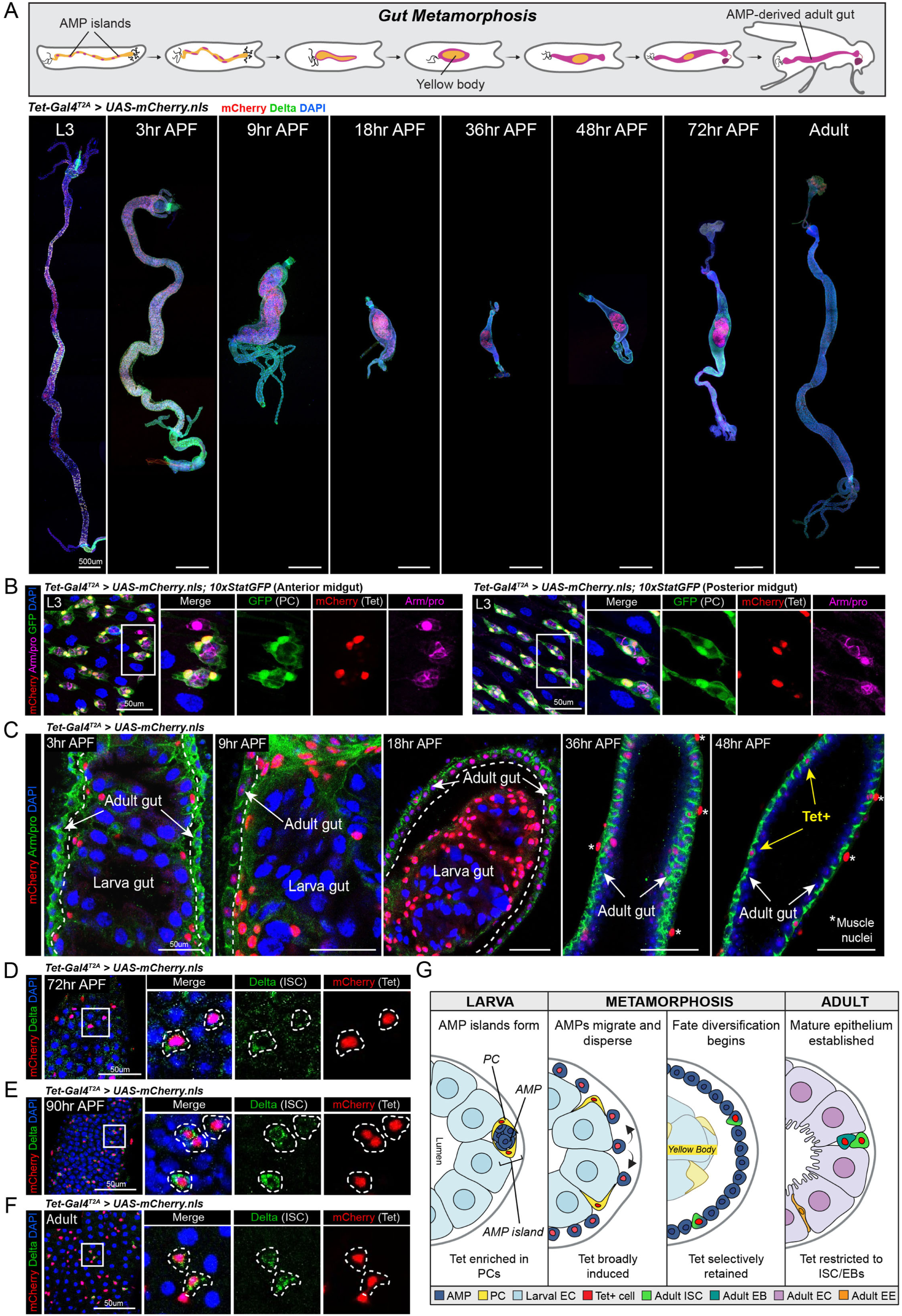
*Tet* expression dynamically marks progenitors during midgut metamorphosis. (A) Overview of gut metamorphosis paired with representative images of *Tet-Gal4^T2A^* > *UAS-mCherry.nls* expression from the larval (L3) stage through adulthood. (B) L3 midguts showing *Tet* expression (red) highly enriched in Peripheral Cells (PCs) marked by *10xStatGFP* (green) and largely absent from AMPs. (C) Temporal dynamics of *Tet* expression (red) during early to late metamorphosis (3hr to 48hr APF), showing broad induction followed by selective retention in the developing adult gut. (D-F) High-magnification images at 72hr APF, 90hr APF, and adult stages demonstrating that *Tet* expression strictly co-localizes with the ISC/EBs. (G) Schematic summary model of AMP morphogenesis, fate diversification, and corresponding *Tet* expression dynamics. It is also worth noting that *Tet* is expressed outside the gut epithelium, in the surrounding visceral muscle.

To functionally test this, we performed lineage tracing using the G-TRACE system combined with inducible *Gal80^ts^* (Evans et al., 2009) and dissected d1 adult guts. Tracing induced at 72hr APF (the stage when *Tet* is highly enriched in Delta+ cells) revealed that *Tet*-retaining progenitors directly gave rise to the adult ISC lineage (Fig. 3). Furthermore, the labeled progeny formed localized multicellular clusters containing both ISCs/EBs and differentiated enterocytes (ECs). The presence of these mature ECs aligns with the established developmental timeline in which ISCs initiate active EC differentiation shortly after eclosion (Z. Guo & Ohlstein, 2015; Takashima et al., 2016). Together, these findings support a model in which *Tet* is initially induced broadly during epithelial remodeling but is subsequently maintained only in progenitors that consolidate ISC identity.

**Figure 3.**
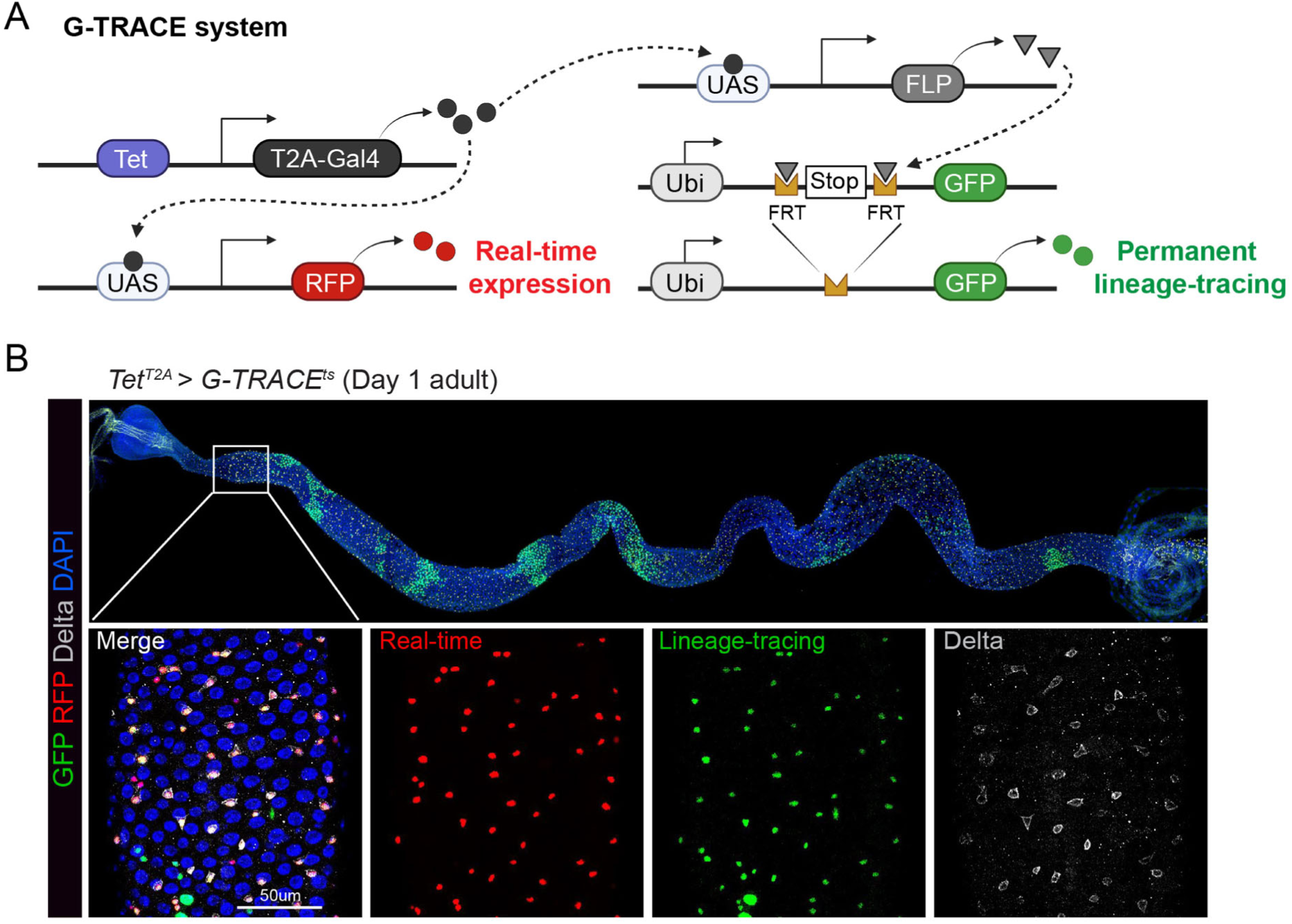
*Tet*-expressing progenitors establish the adult ISC lineage. (A) Schematic representation of the G-TRACE lineage tracing system. (B) Lineage tracing analysis of day 1 adult midguts using Tet-Gal4T2A > G-TRACEts to induce tracing starting from 72hr APF. Real-time Tet expression is marked by RFP (red), and the permanent lineage of Tet-expressing cells is marked by GFP (green), showing strong co-localization with the ISC marker Delta (gray). Note that there are multi-cell clones in different regions of the midgut due to continuous division and differentiation from late pupa stage to the adult.

### Tet function is specifically required during metamorphosis for adult ISC establishment

To determine when Tet is required for adult ISC formation, we performed temporal knockdown experiments using the inducible *esg-Gal4, UAS-mGFP; tub-Gal80ts* system (all timepoints standardized to the equivalent 25°C developmental rate; see Methods). *Tet* depletion restricted to embryonic and larval stages did not impact the adult ISC pool. In contrast, *Tet* knockdown strictly during the pupal stage caused severe ISC loss, demonstrating that Tet is dispensable during larval growth but essential during metamorphosis (Fig. 4A, B).

**Figure 4.**
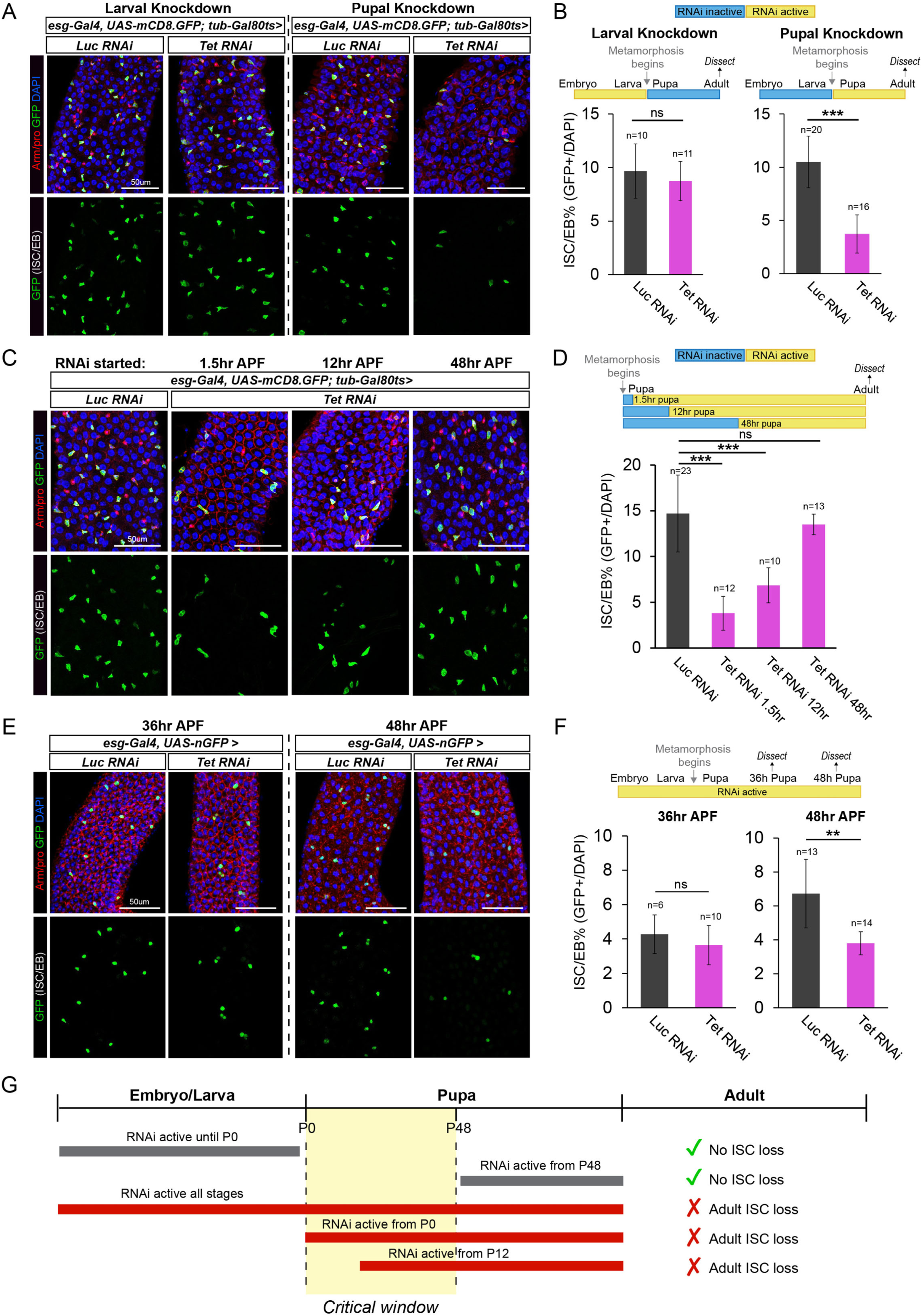
**Tet is required during a specific metamorphic window for adult ISC establishment.** (A) Confocal images of adult midguts following temporally controlled *Tet* knockdown using the *esg-Gal4, tub-Gal80^ts^* system. Knockdown restricted to larval stages does not affect adult ISCs, whereas pupal-restricted knockdown recapitulates the ISC loss phenotype. (B) Quantification of ISC/EB percentage for larval versus pupal knockdown. (C) Midgut images from adults where RNAi was induced starting at 1.5hr, 12hr, or 48hr APF. (D) Quantification of ISC/EB percentage for specific APF RNAi induction timepoints. (E) Midguts dissected at 36hr and 48hr APF following continuous *Tet RNAi*. (F) Quantification of ISC/EB percentage at 36hr and 48hr APF. (G) Summary timeline demonstrating the critical developmental window during the first half of metamorphosis where *Tet* function is required to prevent adult ISC loss. All timepoints are normalized to 25°C developmental timeline (see Methods). Bars represent mean ± SD. *p ≤ 0.05, **p ≤ 0.01, ***p ≤ 0.001. p values were from unpaired Student’s t-test.

To further define this critical window, we induced *Tet* loss at specific intervals during metamorphosis. We found that Tet function is most critically required during early to mid-pupation (0-48hr APF), as RNAi activation from 48hr APF onward did not cause adult ISC loss (Fig. 4C, D). This timing is consistent with our observation that ISC loss is observed as early as 48hr APF in *esg-Gal4>Tet RNAi* guts (Fig. 4E, F). Together, our temporal tracking (summarized in Fig. 4G) reveals that the first 48 hours of metamorphosis constitute the critical time window for Tet activity to ensure proper adult ISC establishment.

### Tet is required for long-term maintenance of stem cell identity in adults

To determine whether Tet is required in established adult ISCs, we performed adult-specific depletion using the inducible *esg-Gal4, UAS-mGFP; tub-Gal80^ts^* system. Adult-specific loss of *Tet* did not acutely disrupt gut structure but instead caused progressive defects over time (Fig. 5A, B). At 10 days post-induction, ISC/EB populations and gut morphology remained normal. By 40 days, however, *Tet*-depleted guts exhibited a marked reduction in ISCs/EBs, accompanied by the appearance of enlarged, polyploid EC-like cells specifically in the anterior midgut. Similar to the developmental loss of *Tet*, the posterior region exhibited no obvious morphological defects (Fig. S3). This enlarged EC phenotype is consistent with compensatory hypertrophy, a well-established response to stem cell depletion where differentiated cells endoreplicate to maintain epithelial integrity (Tamori & Deng, 2013; Z. Guo et al., 2016; Xiang et al., 2017). Together, these findings indicate that Tet is required to sustain stem cell identity in adulthood and that, in its absence, progenitors progressively lose self-renewal ability, leading to gradual stem cell depletion and inducing compensatory EC hypertrophy.

**Figure 5.**
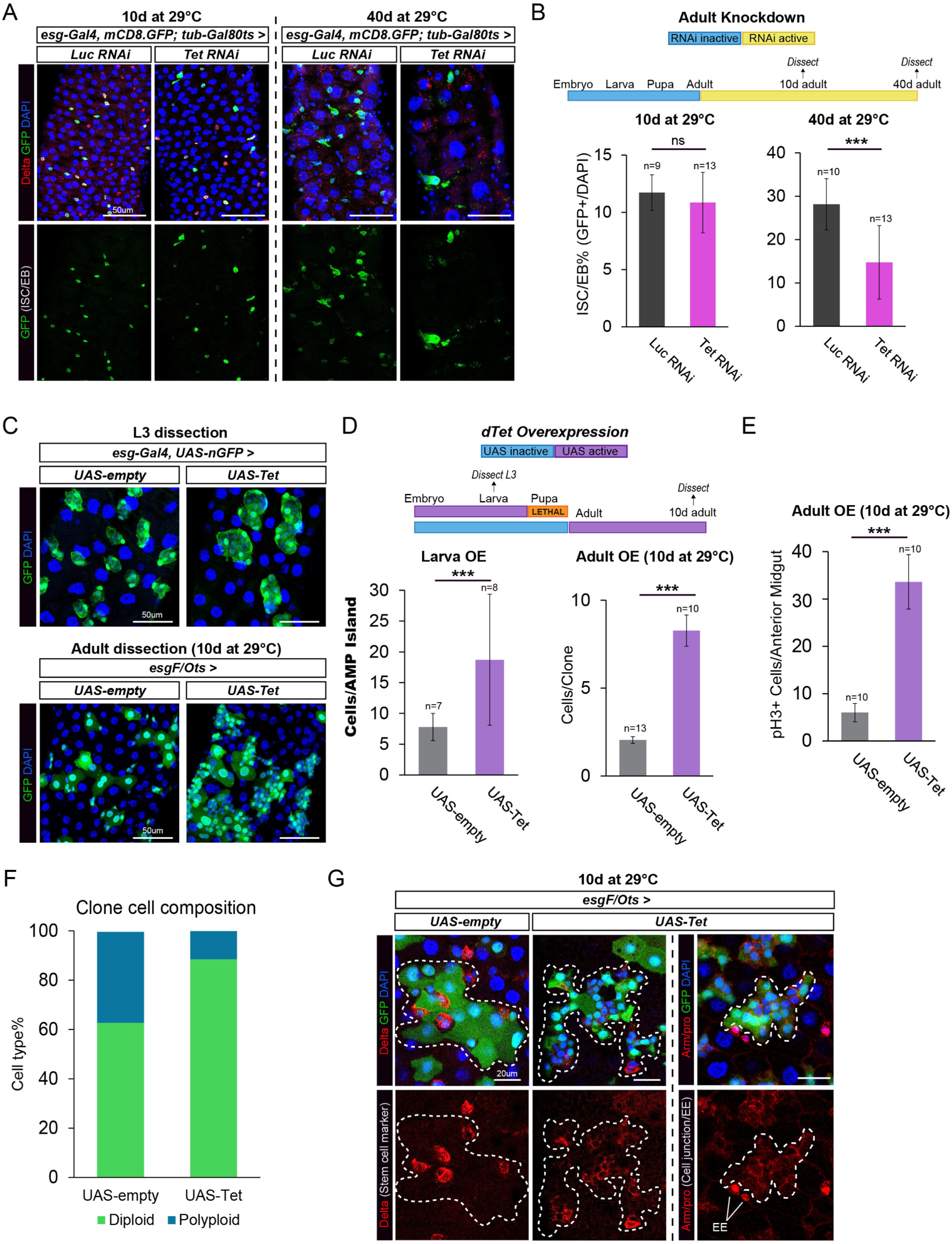
**Tet regulates long-term stem cell maintenance and biases progenitor fate.** (A) Adult midguts subjected to adult-specific *Tet* knockdown (*esg-Gal4, tub-Gal80^ts^*) for 10 days and 40 days at 29°C. (B) Quantification of ISC/EB percentage at 10 and 40 days post-induction. (C) Effects of *Tet* overexpression (OE) in L3 AMP islands (*esg-Gal4*) and in adult midguts after 10 days of induction (*esgF/O^ts^*). (D) Quantification of cells per AMP island in L3 and cells per clone in adult OE midguts. (E) Quantification of proliferating cells (pH3+) per anterior midgut following 10 days of adult OE. (F) Quantification of clone composition (small diploid cells vs large polyploid cells). (G) High-magnification images of clone composition in adult midguts following *Tet* OE, co-stained with the stem cell marker Delta and EE marker Prospero. Bars represent mean ± SD. \**p* ≤ 0.05, \*\**p* ≤ 0.01, \*\*\**p* ≤ 0.001. *p* values were from unpaired Student’s t-test.

### *Tet* overexpression globally biases progenitor fate toward stem cell identity

To further investigate the role of Tet in stem cell fate regulation, we examined the effects of constitutive *Tet* overexpression in progenitors during development using *esg-Gal4*. This was lethal, and we found development was arrested by 48hr APF in *Tet*-overexpressing pupa (Fig. S4A). Thus, precise regulation of endogenous *Tet* levels is essential for successful metamorphosis.

Because this lethality precluded adult midgut analysis, we examined L3 larval progenitors to assess the immediate cellular consequences of elevated *Tet*. This revealed a significant increase in the number of cells within AMP islands in both the anterior (Fig. 5C, D) and the posterior (Fig. S4B, C) compartments, demonstrating a global increase of the progenitor pool. These data suggest that *Tet* overexpression is sufficient to drive progenitor expansion.

To bypass developmental lethality and assess the effect of elevated *Tet* on adult homeostasis, we performed adult-specific overexpression using the temperature-inducible *esgF/O^ts^* system (Jiang et al., 2009). Consistent with the larval phenotype, adult overexpression resulted in a significant increase in clone size and progenitor proliferation (Fig. 5C-E). These expanded *Tet*-overexpressing clones were overwhelmingly composed of small, diploid cells. Immunostaining confirmed these densely packed cells were predominantly Delta+ ISCs, accompanied by a severe reduction in polyploid ECs (Fig. 5F, G). Furthermore, these clusters lacked expression of the enteroendocrine (EE) marker Prospero, confirming they are not EE cells (Fig. 5G). Together, these findings indicate that elevated *Tet* strongly promotes ISC self-renewal and actively suppresses differentiation, reinforcing its role in stabilizing stem cell fate.

### Single-cell transcriptomic atlas of the developing gut

To explore how Tet regulates stem cell establishment, we performed single-nucleus RNA sequencing (snRNA-seq) on control (*Luciferase RNA*i) and *Tet RNAi* midguts across five developmental stages spanning ISC maturation: L3 wandering larva, 0hr white prepupa (P0), 18hr pupa (P18), 48hr pupa (P48), and day 2 adults (D2) (Fig. 6A). Analysis of over 121,000 high-quality nuclei resolved all major gut and gut-associated cell populations, including AMPs, PCs, ISCs/EBs, ECs, EEs, visceral muscle, yellow body, proventriculus (PV) and trachea (Fig. 6B, D). Stage-specific annotations further refined these broad classifications. For example, the larval PV was resolved into six distinct populations at the L3 and P0 stages (detailed annotations and quality control metrics in Fig. S5). Overall, this represents the first single-nucleus transcriptomic atlas of the developing midgut across multiple stages of ISC emergence and provides a comprehensive, temporally resolved resource for studying gut development and tissue remodeling. All data can be visualized, downloaded, and analyzed at https://hongjielilab.org/gut-atlas.

**Figure 6.**
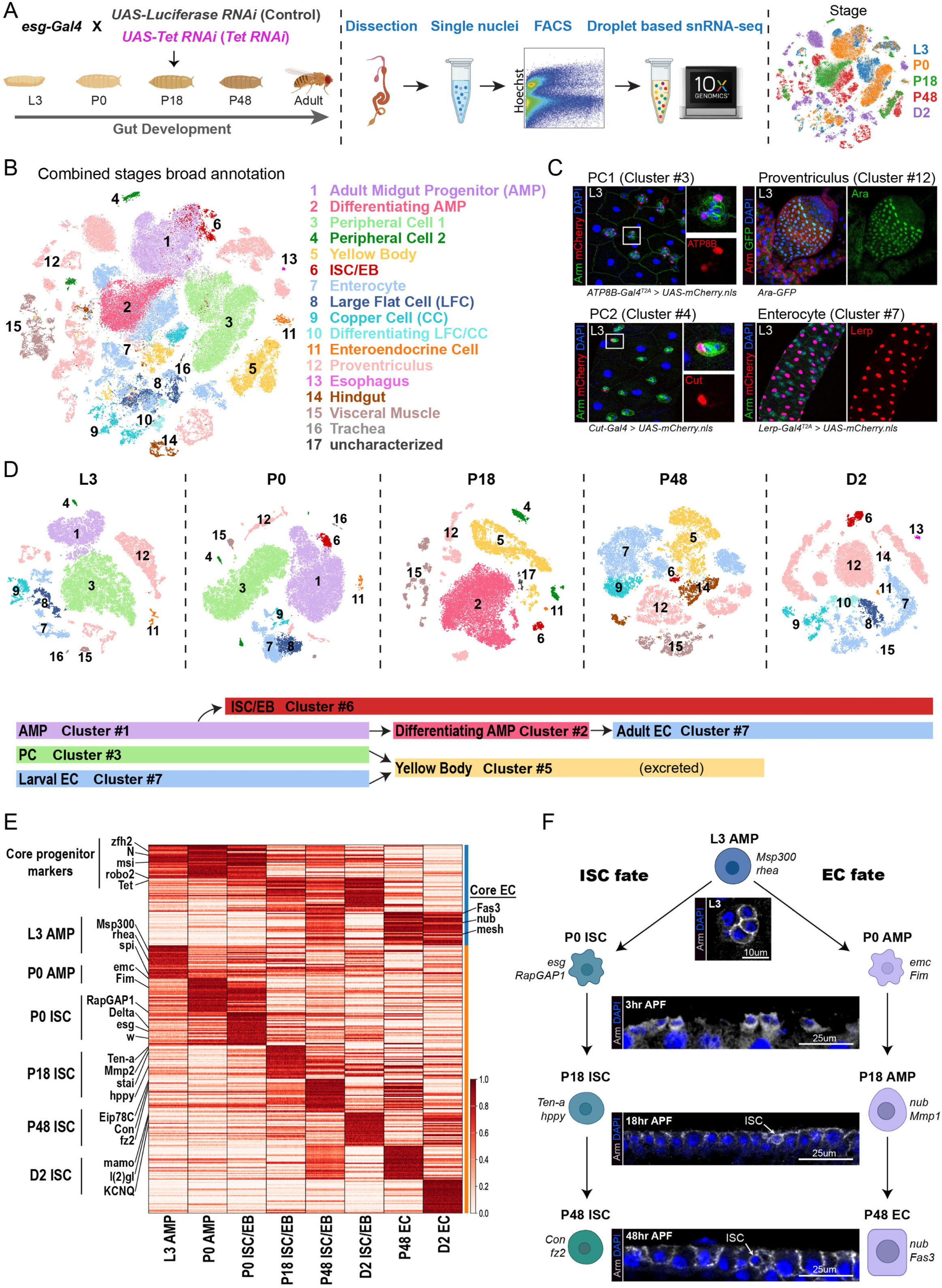
**Single-nucleus transcriptomic atlas resolves the normal maturation trajectory of intestinal progenitors.** (A) Schematic of the experimental design for snRNA-seq spanning five developmental stages covering ISC maturation: L3 larva, 0hr prepupa (P0), 18hr pupa (P18), 48hr pupa (P48), and day 2 adult (D2). (B) Global tSNE projection of over 121,000 high-quality nuclei, resolving all major midgut and gut-associated lineages integrated across development. (C) Validation of L3 stage-specific clusters, identifying unique markers for divergent PC populations (*ATP8B*+ vs *Cut*+), larval ECs (*Lerp+*), and a subset of larval PV cells (*Ara*+). (D) Stage-specific tSNE demonstrating fate divergence beginning at P0. Note that a clear ISC cluster branches out from the AMP population from P0. (E) Heatmap expression profiles tracking the dynamic expression of core lineage-defining genes across the five conceptual phases of ISC maturation in control samples. Side bar color code: blue = broad annotation comparison, orange = stage-specific annotation comparison. (F) Schematic model summarizing the morphological and transcriptomic progression of normal ISC maturation, highlighting the critical bifurcation at P0 into ISC-fated cells (expressing *esg* and *RapGAP1*) and remodeling AMPs (expressing *emc* and *Fim*).

To strengthen our annotation, we performed a series of genetic experiments to determine cluster identity based on cluster-specific genes. As the adult gut has been sequenced in several papers and is well annotated (Dutta et al., 2015; Li et al., 2022; Lu et al., 2023; Parasram et al., 2024; Zhu et al., 2024; Park et al., 2025), we focused these *in vivo* validations on the L3 stage, which has not been thoroughly characterized. Of note, we confirmed two populations of PC cells, a large cluster marked by expression of *ATP8B* (PC1) and a small cluster marked by expression of *Cut* (PC2), which likely represent divergent functions of the two PC populations (Fig. 6C). Additionally, we identified *Ara* as a marker for a subpopulation of the larval PV, and *Lerp* as a marker for larval ECs (Fig. 6C).

Next, we leveraged this temporal resolution to determine when the AMP pool segregates into distinct lineages. Strikingly, our stage-specific analysis revealed that this fate divergence begins at the onset of metamorphosis (Fig. 6D). Progenitors destined to become the future adult ISC/EB population could be transcriptionally distinguished at P0 based on expression of canonical stem cell regulators including *esg*, *Delta*, *Sox100B*, *Sox21a*, and *zfh2*, as well as progenitor competence factors such as *hdc* and the Ras pathway regulator *rau* (Micchelli & Perrimon, 2006; Ohlstein & Spradling, 2006; Resende et al., 2017; Rojas Villa et al., 2019; Meng et al., 2020). The activation of these networks indicates that the transcriptional foundations of ISC identity are established at the very onset of metamorphosis. By P18, these progenitors formed a transcriptionally distinct cluster. Crucially, while *Tet* expression remains initially low in P0 AMPs, it becomes highly upregulated in this distinct progenitor population by P18 (Fig. S6A). This temporal delay indicates that Tet functions not as the initial trigger for divergence, but rather as a critical maturation factor required to safeguard the emerging stem cell lineage.

### Transcriptional programs define the development of intestinal epithelial lineages

To define the transcriptional networks governing lineage maturation, we first identified genes that broadly distinguish progenitors from differentiated ECs across developmental stages (Table S1). Progenitor populations were defined by a coordinated network of transcriptional, signaling, and post-transcriptional regulators (Fig. 6E). This included the transcription factor *zfh2*, signaling receptors such as *Notch* (*N*), *robo2*, and *Egfr*, and RNA-binding proteins like *msi*, *brat*, and *Rbfox1*. Notably, *Tet* was also expressed within this core progenitor network. In contrast, ECs were characterized by activation of a distinct epithelial differentiation program (Fig. 6E). This included genes associated with epithelial structure, adhesion, and metabolic specialization, including *Fas3*, *mesh*, *spir*, *nub*, and *AANAT1*. Together, these transcriptional signatures reflect the acquisition of epithelial barrier function and metabolic activity characteristic of mature ECs.

While these lineage-defining markers establish a broad distinction between multipotent progenitors and terminally differentiated ECs, our temporal dataset allowed us to track the dynamic maturation of the progenitor pool itself. By following this progression across metamorphosis, we found that ISC formation resolves into a continuous developmental sequence organized into five distinct stages:

#### Stage 1: Competence (L3 AMPs exist in a primed progenitor state)

In the larval midgut, AMPs form a relatively homogenous population characterized by high developmental competence. *In vivo* imaging reveals these quiescent cells maintain a round, tightly constrained morphology (L3 in Fig. 6F). These cells do not yet express strong ISC-specific programs (L3 AMP in Fig. 6E). Instead, their gene expression profile reflects high plasticity and responsiveness to future developmental cues. Among the most enriched genes are the cytoskeletal anchor *Msp300* and the core integrin adaptor *rhea* (*Talin*), both of which are maintained throughout subsequent ISC development. They also broadly express the established AMP marker *spi* (*spitz*) (Jiang & Edgar, 2009). This transcriptional landscape indicates that L3 AMPs are maintained in a poised state, primed to respond rapidly to metamorphic signals.

#### Stage 2: Specification (P0 AMPs diverge into ISC-fated and remodeling progenitor populations)

At the onset of metamorphosis (P0), the progenitor pool physically breaks away from their constrained larval niche. They rapidly transition from their round baseline into a highly motile state characterized by dynamic, spiky projections (P0-P3; see Fig. 2C). During this dramatic physical transition, all AMPs strongly upregulate the core adhesion molecule *shg*, likely to maintain tissue cohesion. At the same time, the progenitor pool undergoes a major transcriptional shift, bifurcating into two distinct populations with divergent trajectories (Fig. 6E, F). One group of AMPs activates a stem cell specification program marked by the upregulation of *esg*, *white* (*w*), *Delta*, and *RapGAP1* (Fig. 6E, F; see Fig. 6D P0 cluster #6). In parallel, a second, larger population of AMPs adopts an alternative progenitor state (see Fig. 6D P0 cluster #1). Importantly, these cells remain progenitors and do not immediately differentiate, but instead are transcriptionally reprogrammed to support epithelial growth and remodeling. This population is defined by the upregulation of *emc*, suggesting an active repression of stem cell identity, and factors driving active cytoskeletal dynamics, such as the actin-crosslinking protein *Fim* (Fig. 6E, F).

### Stage 3: Adaptation (P18 ISCs and differentiation-fated AMPs coordinate structural packing while diverging in functional capacity)

Morphologically, this phase is marked by the dampening of early motility. The migratory processes retract to yield a smooth, rectangular structure by P9, before the cells physically pack together to adopt a uniform, rounded geometry by P18 (see Fig. 2C). This structural packing is a collective behavior shared by both the ISC-fated and remodeling progenitor populations as they build the adult epithelium (Fig. 6E, F). Despite this shared physical behavior, their genetic programs diverge to execute distinct roles. The ISC-fated population activates gene programs that enable interaction with and modification of the developing tissue. The strong expression of *Mmp2* and *Ten-a* drives extracellular matrix remodeling and the cell adhesion required for this tight packing, while the microtubule regulator *stai* and the vesicle-trafficking factor *unc-13* facilitate the active cytoskeletal dynamics needed to transition into this rounded cuboidal shape. Alongside these structural changes, the induction of *hppy* activates specialized kinase cascades that integrate nutrient and stress-responsive cues. Meanwhile, the larger remodeling AMP population is fated for differentiation and begins expressing the EC marker gene *nub* and the additional metalloproteinase *Mmp1* (Fig. 6E).

### Stage 4: Integration (P48 ISCs establish structural and signaling connectivity within the developing epithelium)

By mid-metamorphosis (P48), the tissue enters an integration phase where ISCs become physically embedded within the developing adult epithelium. Concurrently, the differentiation-fated AMPs undergo distinct morphological changes. Their overall cell size and nuclei begin to enlarge, making them physically resemble mature ECs rather than early-stage AMPs (Fig. 6E, F). Transcriptionally, the ISC program shifts toward establishing stable cell-cell interactions and coordinated signaling interfaces (Fig. 6E). This transition is marked by the upregulation of adhesion and guidance molecules, including *Sema1a* and *Con* (*Connectin*), alongside the extracellular matrix-interacting factor *pxb*. In parallel, ISCs assemble long-term signaling machinery by expressing the Wnt/Wingless receptor *fz2* and the ecdysone-responsive regulator *Eip78C*. Meanwhile, the differentiation-fated AMPs continue expressing *nub* while upregulating the mature EC marker *Fas3*. Together, these coordinated morphological and transcriptional changes finalize the architecture of the adult gut, with ISCs firmly anchored within their niches and differentiating AMPs maturing into a functional epithelial barrier.

### Stage 5: Stabilization (Adult ISCs acquire homeostatic function and long-term identity maintenance)

By adulthood (day 2), ISCs reach a steady-state identity within the fully formed gut. Transcriptionally, adult ISCs maintain expression of the canonical markers *Delta* and *esg*, while upregulating the transcription factor *mamo*, the potassium ion channel *KCNQ*, and the core apical-basal polarity regulator *l(2)gl* (Fig. 6E).

Collectively, these findings demonstrate that ISC formation is a multi-stage developmental process. Progenitors sequentially acquire competence, undergo lineage specification, adapt to a remodeling environment, integrate into the epithelial niche, and ultimately stabilize as long-lived adult stem cells. All broad and stage-specific markers are summarized in Table S1.

### Tet maintains coordinated transcriptional balance across structural, signaling, and fate-specification programs

Having established a staged model of normal ISC development, we next examined how loss of Tet impacts progression through these phases. tSNE visualization of control and *Tet RNAi* midguts confirms broad capture of the gut lineages (Fig. 7A), while targeted quantification of the ISC/EB pool reveals a clear reduction of adult ISC and EB populations in *Tet*-depleted conditions (Fig. 7B). However, because tSNE dimensionality reduction can fragment continuous lineage progressions, we analyzed the isolated ISCs/EBs using UMAP to accurately preserve their developmental topology. Visualized across this developmental space, *Tet*-depleted progenitors display an obvious transcriptomic shift, forming aberrant clusters that are segregated from control ISC/EB clusters (Fig. 7C). This shift confirms that Tet is required for proper establishment of the adult stem cell lineage.

**Figure 7.**
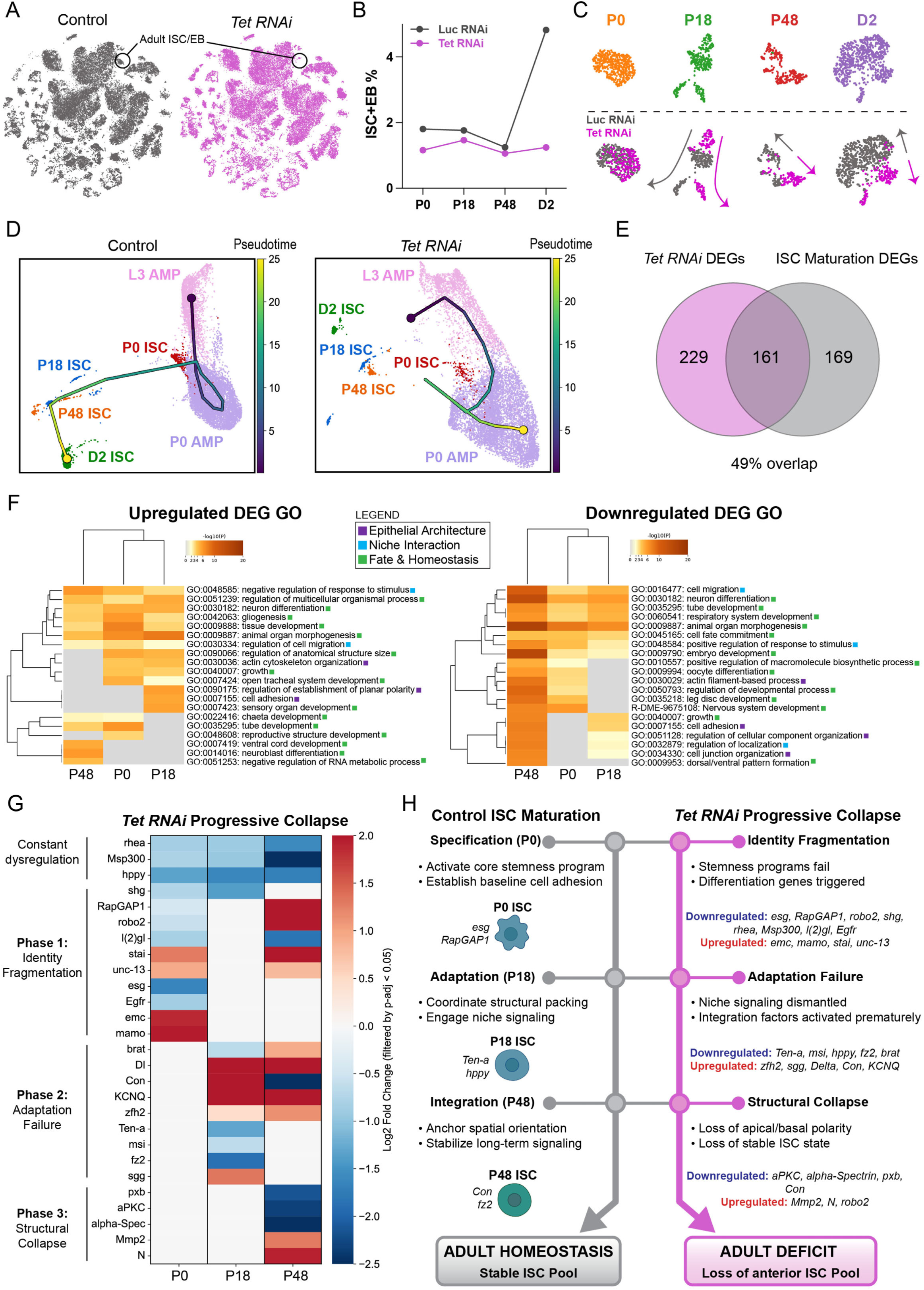
*Tet* loss drives a progressive collapse of the adult ISC pool through temporal and structural dysregulation. (A) Genotype-specific combined stage t-SNE plots. Adult ISC/EB populations are outlined. (B) Quantification of ISC+EB percentages across developmental stages in control versus *Tet RNAi* midguts. Percentage was calculated by dividing the ISC+EB cell number over all midgut epithelial lineage cells. (C) UMAP projection of isolated progenitors (ISC/EBs) from each stage, demonstrating the distinct transcriptomic shift and aberrant clustering of *Tet*-depleted cells. (D) Pseudotime trajectory analysis of control and *Tet RNAi* samples, illustrating the normal continuous progression from L3 AMPs to adult ISCs alongside the severe developmental arrest observed in *Tet RNAi*. (E) Venn diagram highlighting the massive (49%) overlap of two sets of genes: differentially expressed genes (DEGs) between the control and *Tet RNAi*, and stage-specific DEGs underlying normal ISC maturation. (F) Gene Ontology (GO) enrichment analysis of up- and down-regulated *Tet RNAi* DEGs. Functional pathways are assigned to three core categories: epithelial architecture (purple), niche interaction (blue), and fate and homeostasis (green). (G) Temporal heatmap of *Tet RNAi* DEGs within the developing ISC/EB population across metamorphosis, revealing a stepwise deterioration of normal maturation networks. (H) Summary model comparing the wild-type ISC maturation trajectory to the multi-phase progressive collapse induced by *Tet* depletion.

To capture the developmental dynamics of this stem cell loss, we performed pseudotime trajectory inference comparing the progression of control and *Tet*-depleted progenitors (Fig. 7D). In control midguts, progenitors form a continuous developmental trajectory. In contrast, the *Tet RNAi* trajectory exhibits a pronounced developmental arrest. While the early ISC progenitors successfully emerge from the AMP pool, the trajectory subsequently fragments and fails to consolidate into a mature adult ISC population. This directly visualizes the progressive collapse of progenitor development during metamorphosis.

To determine the transcriptomic networks driving this collapse, we compared the *Tet RNAi* differentially expressed genes (DEGs) with genes underlying L3-to-adult developmental ISC maturation. We found a 49% overlap between the two datasets (Fig. 7E; see Table S1). This intersection demonstrates that the transcriptomic dysregulation in *Tet RNAi* largely disrupts normal metamorphic maturation.

To parse the specific functional pathways within this dysregulated network, we performed Gene Ontology (GO) enrichment analysis. The dysregulated genes broadly fell into three major functional categories essential for ISC maturation (Fig. 7F). The first category, epithelial architecture, included specific GO terms such as cell adhesion, actin cytoskeleton organization, and planar polarity establishment. The second category, niche interaction, featured terms covering the regulation of stimulus responses, cell migration, and localization. The third category, fate and homeostasis, captured terms involving tissue development, cell fate commitment, and cell differentiation. Furthermore, these functional categories were enriched in both upregulated and downregulated gene sets. This indicates that Tet regulates a complex transcriptional balance rather than functioning as a simple binary switch.

### *Tet* loss drives a progressive collapse of the adult ISC pool through temporal and structural dysregulation

Temporal DEG analysis revealed that *Tet* depletion does not trigger immediate progenitor loss, but instead induces a stepwise deterioration of the ISC maturation program. This collapse can be resolved into three mechanistically distinct phases, characterized by the fragmentation of core stem cell identity and a severe temporal mismatch that drives the out-of-phase activation of normal maturation milestones (Fig. 7G, H).

### Phase 1: P0 (Identity fragmentation and fate mis-specification)

At P0, *Tet RNAi* ISCs successfully bifurcate from the AMP pool but fail to establish a coherent stem cell program. Essential P0 maturation markers are significantly reduced, including the stemness factor *esg*, the signaling regulators *RapGAP1* and *robo2*, and niche-sensing components like *Egfr*. This incomplete specification is coupled with severe fate misregulation. Progenitors inappropriately upregulate *emc*, normally restricted to differentiating AMPs, while prematurely activating the adult ISC transcription factor *mamo*. Concurrent failure to maintain baseline structural and adhesion molecules (*shg*, *rhea*, *Msp300*) is coupled with downregulation of the polarity regulator *l(2)gl* and the kinase *hppy*, further destabilizing identity. Paradoxically, these progenitors also exhibit an out-of-phase upregulation of dynamic cytoskeletal and trafficking factors, such as *stai* and *unc-13*. Consequently, rather than locking in a stable identity, *Tet*-depleted ISCs exhibit a conflicting transcriptional profile characterized by out-of-phase differentiation signals.

#### Phase 2: P18 (Adaptation failure and loss of asymmetric fate control)

As control cells enter the P18 adaptation phase, *Tet RNAi* ISCs fail to deploy the networks required for structural packing and functional adaptation. Essential adhesion and RNA-regulatory factors (*Ten-a*, *msi*) are strongly downregulated, with the sustained loss of core structural anchors (*shg*, *rhea*, *Msp300*). Beyond this structural gene disruption, signaling coordination is actively dismantled. This is evidenced by the reduction of the Wnt receptor *fz2* and the concurrent elevation of its antagonist, *sgg*. In parallel, asymmetric fate segregation is compromised. The fate determinant *brat* is downregulated, while the Notch ligand *Delta* sharply increases. Furthermore, *Tet*-depleted ISCs prematurely activate the integration factor *Con* and the stabilization marker *KCNQ* while simultaneously overexpressing *zfh2*. Ultimately, the loss of structural anchors and the severe uncoupling of fate determinants indicate an inability to sustain coordinated adaptation.

#### Phase 3: P48 (Niche integration failure and structural collapse)

By mid-metamorphosis (P48), these defects culminate in failed niche integration. *Con*, having aberrantly peaked at P18, is depleted when required for anchoring, and the integration factor *pxb* is similarly disrupted. Epithelial architecture collapses as polarity scaffolds essential for division are downregulated, including *aPKC* and *alpha-Spectrin*. This occurs alongside the continuous depletion of *l(2)gl*, *rhea*, and *Msp300*. At the same time, aberrantly elevated *Mmp2* promotes the degradation of basement membrane attachments. Moreover, it shows an increase of of *robo2* and *zfh2*, sustained expression of *KCNQ*, and an out-of-phase upregulation of previously suppressed factors (*RapGAP1*, *brat*, *stai*, *unc-13*). Then a surge in the Notch receptor (*N*) emerges. Driven by sustained *Delta*, this intense Notch signaling likely forces the ISCs to bypass the transit-amplifying EB state. *In vivo* analysis confirmed that *NRE-lacZ-*positive EB populations do not accumulate (Fig. S6B, C), supporting a model where ISCs fail to maintain a stable intermediate and instead undergo rapid, Notch-driven differentiation into ECs. This is consistent with the observation that high Notch activity drives ISC for EC differentiation instead of for stem cell renewal or maintenance (Kapuria et al., 2012).

Collectively, this stepwise progression reveals that Tet functions as a central transcriptional regulator of progenitor identity. To determine if a single downstream target could account for this collapse, we performed a targeted genetic screen of 51 fly lines for 25 top misregulated genes, such as *Ten-a*, *Msp300*, *fz2*, *hppy*, *Delta*, and *KCNQ*. This screen revealed that single-gene manipulation is not sufficient to phenocopy the anterior ISC loss observed in *Tet RNAi* (Table S2). This confirms that Tet is required to stabilize a broad and highly integrated regulatory network rather than acting through a single dominant effector.

## DISCUSSION

Here, we identify Tet as a critical intrinsic regulator of stem cell fate required for ISC establishment during *Drosophila* gut development. Tet coordinates a network of genes spanning epithelial integrity, niche signaling, and fate maintenance. It acts as an essential transcriptional safeguard, maintaining the precise regulatory balance required to stabilize stem cell programs against premature differentiation. Without Tet, ISCs lose the structural anchors and polarity machinery necessary for asymmetric division and fail to correctly integrate stabilizing Wnt and Notch cues. This multi-layered regulatory collapse depletes the stem cell pool before adulthood, severely compromising the regenerative capacity required for long-term homeostasis.

### Regional intestinal stem cell specification and the threshold of signaling competence

The distinct regional vulnerability of the anterior midgut following *Tet* depletion aligns with established models of midgut compartmentalization. Although *Tet* is broadly expressed in ISCs across the midgut, only anterior progenitors fail to survive. This regional sensitivity indicates that anterior progenitors likely operate closer to a critical developmental threshold and therefore depend more strongly on intrinsic transcriptional regulators to navigate metamorphosis.

Recent work demonstrates that the precise integration of Wnt pathway activation and Notch pathway suppression at approximately 12.5hr APF is required for successful ISC specification (Y. Wu et al., 2025). Our transcriptomic data aligns with this critical developmental window. Following *Tet* depletion, progenitors show a collapse of Wnt receptivity by P18, characterized by the loss of the Wnt receptor *fz2* and the upregulation of the Wnt antagonist *sgg* (Fig 7G). These components indicate that wild-type ISCs are actively organizing their membrane architecture to support efficient signal transduction and intercellular communication, a process that completely fails in the absence of Tet. Crucially, this Wnt shutdown occurs alongside a failure to suppress Notch signaling, evidenced by the loss of the Notch-repressive coregulator *spen* and the upregulation of the Notch ligand *Delta*.

This shutdown of Wnt responsiveness and precocious shift toward Notch activation provides a possible mechanistic explanation for the regional phenotype. The *Drosophila* midgut is patterned by regionalized gradients of Wingless (Wg) signaling, with concentrated sources at both the anterior (cardia) and posterior (hindgut) boundaries (Buchon et al., 2013; Tian et al., 2016). Because anterior progenitors develop immediately adjacent to the cardia, we propose they may exhibit a heightened reliance on this local Wnt signal for self-renewal, potentially lacking redundant niche survival pathways present in the posterior midgut. Consequently, these anterior cells may strongly depend on intrinsic transcriptional regulators, such as Tet, to maintain a transcriptional state that maximizes Wnt receptivity (*fz2*) while dampening antagonistic or differentiation-driving pathways (*sgg*, *Notch*). When Tet is lost, the subsequent reduction of Wnt receptors renders the anterior ISCs largely insensitive to the local Wg source, facilitating a premature transition into Notch-driven differentiation. Importantly, this transcriptional threshold model can also explain the global expansion of progenitors observed when *Tet* is overexpressed, as a sustained Wnt-receptive state would likely drive enhanced proliferation across all regional compartments.

Driven by sustained *Delta*, this intense Notch signaling likely forces progenitors to bypass the transit-amplifying EB state. This supports a model where *Tet*-depleted progenitors fail to lock in stemness, cannot maintain a stable intermediate, and instead undergo rapid Notch-driven differentiation into ECs (Kapuria et al., 2012). Together, these findings support a paradigm in which Tet regulates the transcriptional programs required for signaling integration, thereby enabling cells to successfully interpret fate cues.

### Tet preserves progenitor fate by coupling epithelial polarity and niche adhesion

Recent work has demonstrated that during embryogenesis, AMPs originate from the asymmetric division of neuroblast-like endodermal precursors (Plygawko et al., 2025). This transient competence state relies on the coordinated deployment of apical-basal polarity, asymmetric division machinery, and precise niche adhesion. Our transcriptomic analysis suggests that AMPs must reactivate this neuroblast-like program during metamorphosis to establish the adult ISC pool, with *Tet* acting as a key transcriptional regulator of this structural and developmental state.

Rather than simply stalling development, *Tet* depletion disrupts the core architecture of progenitor cells. Loss of *Tet* removes the apical polarity components, including *aPKC* and *cno,* that are required to orient the mitotic spindle, while simultaneously dysregulating basal fate determinants such as *brat* and *numb*. This coordinated breakdown of polarity machinery likely reflects a physical failure of asymmetric division at a stage when the gut epithelium is actively remodeling and experiencing mechanical stress. Because the progenitor pool undergoes a synchronized wave of proliferation at roughly 48hr APF (Jiang & Edgar, 2009), the dismantling of polarity machinery occurs precisely when the capacity for asymmetric division is most critically required. Consistent with this interpretation, Tet has been implicated in maintaining tissue architecture in other contexts, including glial organization in the brain where it interfaces with Hippo signaling (Ismail et al., 2019; Frey et al., 2022), a mechanotransduction pathway that integrates adhesion and polarity to control cell fate (Chen et al., 2010; Cai et al., 2018; Zheng & Pan, 2019).

In addition to polarity, our data indicate that Tet is required to establish basement membrane adhesion complexes that anchor progenitors within their niche, consistent with recent findings in *Drosophila* flight muscle development (Gerdy et al., 2025). We observe widespread dysregulation of the integrin extracellular matrix interface in *Tet*-depleted progenitors, including persistent downregulation of *rhea* (*Talin*) and stage-specific misregulation of *trol* (*Perlecan*) and *Dg* (*Dystroglycan*). These factors collectively support the establishment of stable cellular architecture, enabling ISCs to maintain defined spatial relationships within the epithelial sheet. Because stable niche adhesion is a fundamental determinant of epithelial stem cell identity (Song & Xie, 2002; Morrison & Spradling, 2008; Lin et al., 2013), this loss of structural anchorage severely destabilizes progenitor competence. In the absence of coordinated polarity and adhesion, divisions become biased toward differentiation, ultimately leading to progressive depletion of the ISC pool.

### Conserved roles of Tet proteins in epithelial architecture and stem cell regulation

Our findings are consistent with a broader, evolutionarily conserved relationship between Tet proteins, transcriptional stability, and epithelial integrity. Mammalian Tet enzymes are heavily implicated in regulating epithelial architecture and barrier stability through the control of transcriptional programs governing tissue organization and stem cell behavior. In the mammalian gut, Tet1-mediated hydroxymethylation regulates Wnt signaling and stem cell programs required for homeostatic renewal (Kim et al., 2016). Furthermore, Tet3 promotes intestinal epithelial integrity during stress by supporting protective transcriptional responses (Gonzalez et al., 2023). Similarly, Tet2 is required to maintain epithelial barrier function in airway epithelia (Qin et al., 2020), and integrin-dependent signaling actively interfaces with Tet activity to regulate epithelial junctional organization (Ma et al., 2022).

Collectively, these studies support a conserved role for Tet proteins in stabilizing epithelial architecture and stem cell identity across tissues. Our findings significantly extend this concept by identifying Tet as a key intrinsic regulator of progenitor fate during ISC development and homeostasis. These established gene networks reflect the acquisition of a mature epithelial program that supports long-term ISC function, structural polarity, and steady-state renewal within the adult gut environment. We propose that Tet-mediated stabilization of epithelial architecture, polarity, and niche signaling represents a conserved mechanism that ensures robust stem cell fate specification during development and tissue remodeling.

## RESOURCE AVAILABILITY

### Lead contact

Further information and requests for resources and reagents should be directed to and will be fulfilled by the lead contact, Hongjie Li.

## Materials Availability

This study generated snRNA-seq data which are available as described below.

## ACKNOWLEDGMENTS

We would like to thank Drs. Hugo Bellen, Zheng Zhou, Andre Catic and David Nelson for their knowledge and constructive feedback. We thank Drs. Bing Yao, Peng Jin, Guoqiang Zhang, Dahua Chen, Hugo Bellen, Lucy O’Brien, Henrich Jasper, and Norbert Perrimon for sharing fly lines.

## AUTHOR CONTRIBUTIONS

Conceptualization: N.A., T.P.W., H.L. snRNA-seq experiment: N.A., Y-J.P., Y.Q. snRNA-seq data processing and quality control: Y-J.P snRNA-seq data annotation: N.A., Y-J.P., C-Y.L., T-C.L. snRNA-seq data analysis: Y-J.P., N.A., T.J.

Data interpretation: N.A., Y-J.P., T.J., C-Y.L., T-C.L., S.G., Z.Y., B.S., Y.Z., H.L.

Fly *in vivo* experiments: N.A., S.E., Y.Y., T.J.

Writing: N.A., H.L.

Review and Editing: All authors. Supervision: H.L.

## METHODS

### Fly Stocks

Stocks were maintained on standard cornmeal-yeast-agar medium at 25°C unless otherwise indicated. For RNAi-mediated knockdown of *Tet*, the following lines were used: *Tet RNAi* #1 (VDRC #110549), *Tet RNAi* #2 (from Bing Yao and Peng Jin, Yao et al. 2018), and *Tet RNAi* #3 (VDRC #27098). Additional lines included *UAS*-*Tet* (from Dahua Chen, Zheng et al. 2015), *Tet-Gal4^T2A^* (BDSC #76666), *10xStatGFP* (BDSC #26198), *UAS-redstinger, UAS-Flp, Ubi-FRT.stop.GFP/cyo* (G-TRACE, BDSC #28280), *UAS-Luciferase RNAi* (BDSC #31603), *UAS-empty* (from Hugo Bellen, Moulton et al. 2021), *esgF/O^ts^* (from Bruce Edgar, Jiang et al., 2009), *esg-Gal4, UAS-nGFP* (from Lucy O’Brien), *NRE-LacZ; esg-Gal4, UAS-nGFP* and *esg-Gal4, UAS-GFP; tub-Gal80ts* (from Heinrich Jasper), *esg-Gal4* (from Shigeo Hayashi, Hayashi et al. 2002), *UAS-mCherry.nls* (BDSC #38424), *T2A-ATP8B* (BDSC #91431), *Cut-Gal4* (BDSC #27327), *Ara-GFP* (BDSC #93570), and *T2A-Lerp* (BDSC #77798).

For temporal control of Gal4 activity, flies carrying *tub-Gal80ts* were maintained at 18°C and shifted to 29°C at defined developmental stages as described below.

### Immunofluorescence Staining

Larval, pupal, and adult midguts were dissected in 1X phosphate buffered saline (PBS). For standard immunostaining, tissues were fixed in 4% paraformaldehyde (VWR #100496-494) for 45 minutes at room temperature. For Delta staining, a modified fixation protocol was utilized: midguts were fixed in a mixture of 4% PFA and n-heptane for 15 minutes at room temperature, followed by a brief dehydration in 100% methanol and subsequent stepwise rehydration in PBST (PBS supplemented with 0.1% Triton X-100).

Following their respective fixations, all tissues were washed in PBST and blocked in 5% normal goat serum (BioLegend #927503) in PBST for 1 hour at 4°C. Samples were incubated with primary antibodies diluted in blocking solution overnight at 4°C. After washing in PBST, tissues were incubated with species-appropriate secondary antibodies diluted in blocking solution for either 4 hours at room temperature or overnight at 4°C. Nuclei were counterstained with DAPI (1:1000; Thermo Fisher Scientific #62248) for 20 minutes at 4°C prior to mounting in SlowFade Gold Antifade Mountant (Thermo Fisher Scientific #S36936).

### Antibodies

Primary antibodies used were mouse anti-Delta (Developmental Studies Hybridoma Bank [DSHB], C594.9B; 1:100), mouse anti-Armadillo (DSHB, N2 7A1; 1:100), mouse anti-Prospero (DSHB, MR1A; 1:250), rabbit anti-phospho-Histone H3 (Ser10) (Cell Signaling Technology #9701; 1:1000), and rabbit anti-β-galactosidase (MP Biomedicals #559761; 1:500).

Secondary antibodies included donkey anti-mouse Alexa Fluor 488 (Jackson ImmunoResearch #715-545-151), donkey anti-mouse Cy3 (Jackson ImmunoResearch #715-165-151), donkey anti-mouse Alexa Fluor 647 (Jackson ImmunoResearch #715-605-150), donkey anti-rabbit Alexa Fluor 568 (Thermo Fisher Scientific #A10042), and goat anti-rabbit Alexa Fluor 647 (Jackson ImmunoResearch #111-605-144).

### Oxidative Stress (Paraquat) Survival Assay

To assess susceptibility to oxidative stress, adult female flies of the indicated genotypes were first housed at 25°C and allowed to mate for several days after eclosion until they were 7 days old. Prior to treatment, flies were starved for six hours in empty vials at 25°C to promote consistent and synchronized feeding. Flies were then transferred into empty vials containing a filter paper disc (Whatman) saturated with either a control solution of 5% sucrose or an experimental solution of 5 mM paraquat (Sigma-Aldrich, #856177) dissolved in 5% sucrose. Approximately 15-20 flies were housed per vial, with a minimum of 50 flies analyzed per genotype and condition. Survival was monitored continuously, and mortality rates were recorded at 24 and 48 hours post-exposure. Survival data were analyzed for statistical significance at each time point using a Student’s t-test.

### Lifespan Assay

For lifespan analysis, newly eclosed adults of the indicated genotypes were collected and housed in bottles containing standard cornmeal medium for 2-3 days to allow mating. Females were then separated out into lifespan cages, with approximately 100 flies per genotype. Fly food was replaced every 2-3 days, and the number of dead flies was recorded at each transfer until all flies had died. Survival curves were generated and analyzed using OASIS 2 (Han et al., 2016). Lifespans were performed in duplicate, with the total sample size of control being 240 flies, *Tet RNAi* #1 being 247 flies, and *Tet RNAi* #2 being 118 flies.

### Temporal Induction

To temporally control *Tet* knockdown or overexpression, flies carrying *esg-Gal4* and *tub-Gal80ts* were crossed to *UAS-Tet RNAi* or *UAS-Tet* and maintained at 18°C to suppress Gal4 activity. Progeny were shifted to 29°C at defined developmental stages to induce transgene expression. For larval stage induction, animals were crossed and maintained at 29°C until pupariation, at which point they were shifted to 18°C. For pupal stage induction, animals were maintained at 18°C through larval development and shifted to 29°C at specified pupal stages. For adult-specific induction, 2-3 day old flies were shifted to 29°C and maintained for the duration of the experiment. Control animals were treated identically.

### Developmental Staging and White Prepupa Collection

To ensure highly precise developmental staging for all temporally controlled experiments, wandering third instar (L3) female larvae of the appropriate genotypes were isolated and visually monitored every 30 minutes. The 0hr APF (white prepupa) timepoint was strictly defined as the onset of pupariation, characterized by the complete cessation of larval movement, eversion of the anterior spiracles, and the initial hardening of the white pupal case. All subsequent aging for pupal timepoints (e.g., 1.5hr, 12hr, 36hr, and 48hr APF) was calculated from this strictly defined 0hr APF baseline.

### Normalization of developmental timing to the 25°C timeline

To accurately stage animals during temperature-shift experiments, developmental progression at the 18°C restrictive temperature was normalized to the canonical *Drosophila* developmental timeline at 25°C. Because development at 18°C proceeds at approximately half the rate of development at 25°C, all physical aging intervals prior to the 29°C temperature shift were doubled to reach the desired developmental equivalent. For example, to induce RNAi at a developmental stage equivalent to 12hr APF on the 25°C timeline, white prepupae (0hr APF) were collected and aged at 18°C for 24 hours before being shifted to 29°C. Similarly, for RNAi induction at the 48hr APF equivalent, white prepupae were collected and aged at 18°C for 96 hours prior to the 29°C shift. This normalization ensures that RNAi activation precisely corresponds to the established morphological and transcriptional milestones of the standard 25°C developmental timeline.

## Single-nucleus RNA-seq

### Tissue Preparation

Midgut samples were collected from females across each stage: wandering third instar larva (L3), white prepupa (P0), 18hr pupa (P18), 48hr pupa (P48), and day 2 adults (D2). Guts were dissected in cold Schneider medium. Crop, malpighian tubules, and hindgut were removed to enrich midgut cells during sequencing. Guts were placed into 1.5ml RNase free Eppendorf tubes with a cold Schneider medium containing RNase Inhibitor (Promega, N2611), flash-frozen using liquid nitrogen and stored at -80°C until processing. Sample sizes for each stage are as follows: L3 midguts n=50 per genotype; P0 n=40 per genotype; P18 n=250 for control and n=360 for *Tet RNAi*; P48 n=120 per genotype; D2 n=50 per genotype.

### Nuclei Isolation and FACS

Single-nucleus suspensions were prepared following the protocol described previously (McLaughlin et al., 2022), except that homogenization was limited to 15 strokes with a tight pestle. Next, we used the BD AriaIII FACS sorter to collect nuclei. Nuclei were stained by Hoechst-33342 on ice (1:1000; >5min). Hoechst+ nuclei were collected during sorting. Individual nuclei were collected into one 1.5ml RNase-free Eppendorf tube with 200µl 1x PBS with 0.5% BSA (RNase inhibitor added) as the receiving buffer. For each sample, 100k nuclei were collected. Nuclei were spun down for 10 min at 950g at 4°C and then resuspended using 40µl or desired amount of 1x PBS with 0.5% BSA (RNase inhibitor added). 2µl of nucleus suspension was used for counting the nuclei with hemocytometers to calculate the concentration. 24K nuclei were loaded to the Chromium Next GEM Single Cell 3’ Reagent Kits v3.1 (Dual Index) for each channel to target >10k nuclei.

### Library Preparation and Sequencing

Next, we performed snRNA-seq using the 10x Genomics platform with the Chromium Next GEM Single Cell 3’ Reagent Kits v3.1 (Dual Index) with the following settings. All PCR reactions were performed using the BioRad C1000 Touch Thermal cycler with a 96-deep Well Reaction Module. For cDNA amplification, 12 cycles were performed. Sample index PCR was conducted following the 10x Genomics protocol recommended cycle numbers. As per the 10x protocol, 1:10 dilutions of amplified cDNA and final libraries were evaluated on a bioanalyzer. The final library was sent to Admera Health, LLC for Illumina NovaSeq X plus 25B 2x150 lane sequencing with the dual index configuration Read 1 28 cycles, Index 1 (i7) 10 cycles, Index 2 (i5) 10 cycles, and Read 2 90 cycles. A PhiX control library was spiked in at 5% concentration. The sequencing depth is about 37-56K reads per nucleus.

## Bioinformatics and Computational Analysis

### snRNA-seq data processing

Raw snRNA-seq data, in the form of FASTQ files, underwent alignment to the *Drosophila melanogaster* reference genome (FlyBase release 6.31) using the Cell Ranger software (v8.0.0). Subsequent steps involved the removal of ambient RNA contamination via CellBender (Fleming et al., 2023) and the identification and exclusion of potential doublet cells using Scrublet (Wolock et al., 2019). Quality control criteria necessitated the elimination of cells exhibiting fewer than 200 genes. Genes detected in fewer than 3 nuclei were removed from our analysis. Additionally, cells harboring over 5% of mitochondrial transcripts were eliminated. The majority of the snRNA-seq data analysis was conducted using the Scanpy package (v1.10.3) (Wolf et al., 2018).

### Cell type annotation and clustering

To annotate cell types across ten distinct samples (control and RNAi for L3, 0h APF, 18h APF, 48h APF and D2), we employed a multi-step integration and manual curation strategy. Initial cell type labels were predicted using a logistic regression classifier trained on the Fly Cell Atlas (FCA) adult gut dataset (Li et al., 2022). While this provided a reference for D2 samples, manual annotation was prioritized for developmental stages. We performed Harmony-based integration for each stage to align genotypes (control and RNAi) and generated stage-specific t-SNE and UMAP embeddings. Clusters were defined using the Leiden algorithm at multiple resolutions and manually annotated based on established marker genes, cross-referenced with the transferred labels. For poorly characterized clusters, candidate markers were identified and validated through *in vivo* experiments across developmental stages. Finally, all samples were integrated using Harmony to account for age and genotype, allowing for consistent cell type refinement across the entire dataset.

### Differential gene expression and Gene ontology analysis

To characterize cell-type-specific gene expression profiles, we identified marker genes from the control (*Luc RNA*i) samples across developmental stages and lineages. Using the Wilcoxon rank-sum test within the Scanpy framework, we prioritized 30 broad markers per cell category, followed by the selection of 30 stage-specific markers for each group along the developmental trajectory. Statistical significance was defined by a Benjamini-Hochberg adjusted p-value (false discovery rate, FDR) < 0.05. To ensure specificity, genes already assigned as broad markers were excluded from the stage-specific selection. The resulting expression patterns were visualized via a heatmap to highlight both lineage-conserved and stage-dependent transcriptional signatures. DEGs between genotypes within the ISC/EB clusters were also identified using the Wilcoxon rank-sum test with the same FDR threshold, followed by functional enrichment analysis using Metascape (Zhou et al., 2019).

### Cell type proportion analysis

To evaluate cellular composition across samples, we calculated the relative ratio of the ISC/EB cluster. For each sample, the ratio was determined by dividing the number of ISC/EB nuclei by the total number of midgut-lineage nuclei. To ensure a consistent baseline, we excluded non-midgut contaminating tissues (visceral muscle, hindgut, and trachea) from the total count. However, developmental structures originating from the midgut epithelium, such as the yellow body during pupal stages, were retained in the normalization pool. These ratios were subsequently compared across different developmental stages and genotypes.

### Lineage Trajectory and Pseudotime Analysis

To reconstruct developmental trajectories, we first sub-selected the AMP and ISC/EBs from the integrated dataset. The data was then split by genotype into control and *Tet RNAi* groups. For each group, PAGA was employed to estimate cluster connectivity and generate a coarse-grained topology (Wolf et al., 2019). Subsequently, scFates was used to calculate pseudotime values with the L3 AMP defined as the root (Faure et al., 2023).

## DATA AVAILABILITY

Raw FASTQ files, expression matrix, and processed h5ad files, including celltype annotations, are available at NCBI/GEO (accession number GEO: GSE327108). All annotated data are also available for visualization, download, and custom analysis at https://hongjielilab.org/gut-atlas.

## Supplementary Information

**Figure S1.**
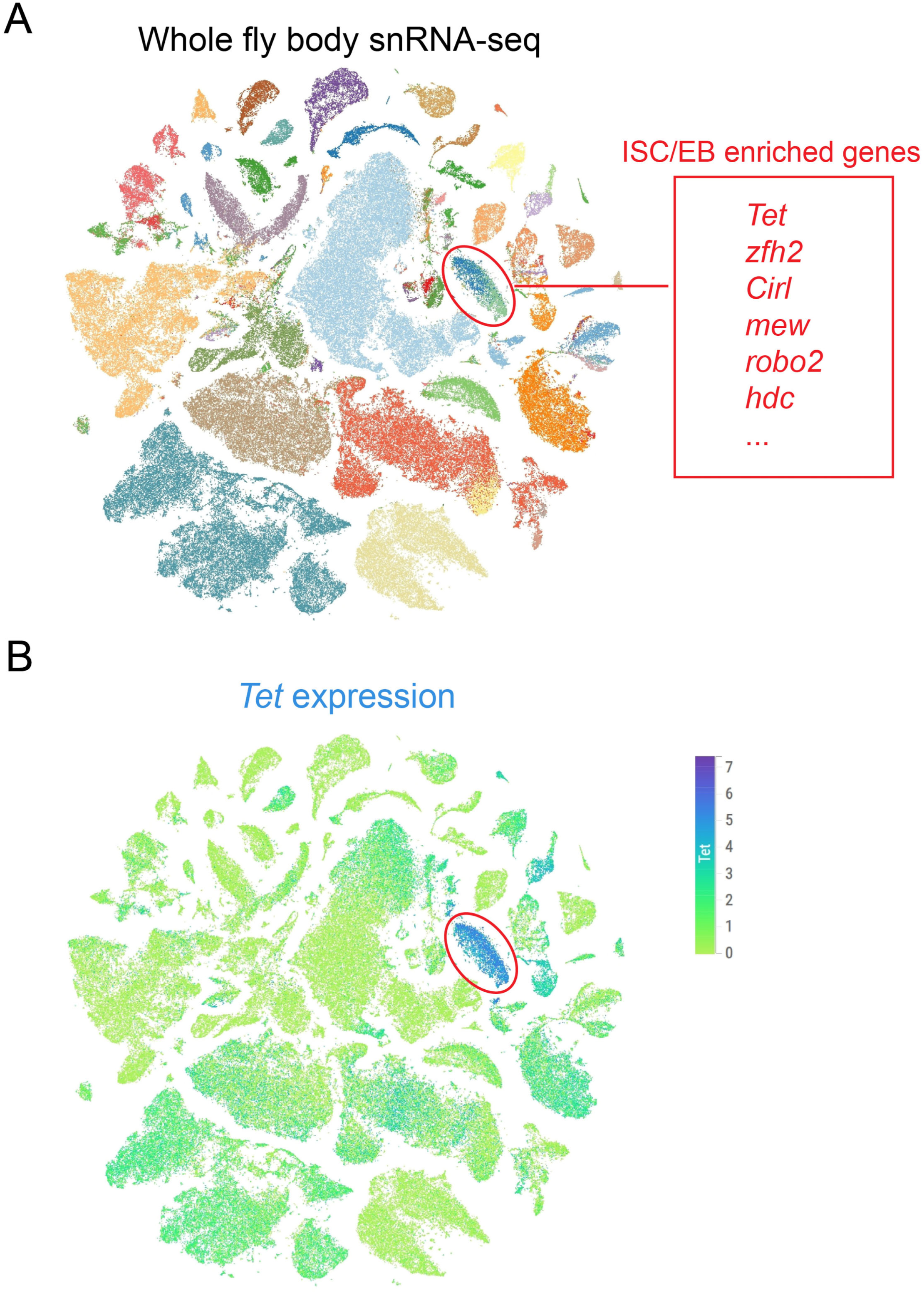
*Tet* is enriched in adult ISC/EBs (A) Whole fly body snRNA-seq tSNE from the AD-FCA highlighting the ISC/EB cluster and listing the top enriched genes. (B) tSNE demonstrating that *Tet* is highly expressed in the ISC/EB cluster.

**Figure S2.**
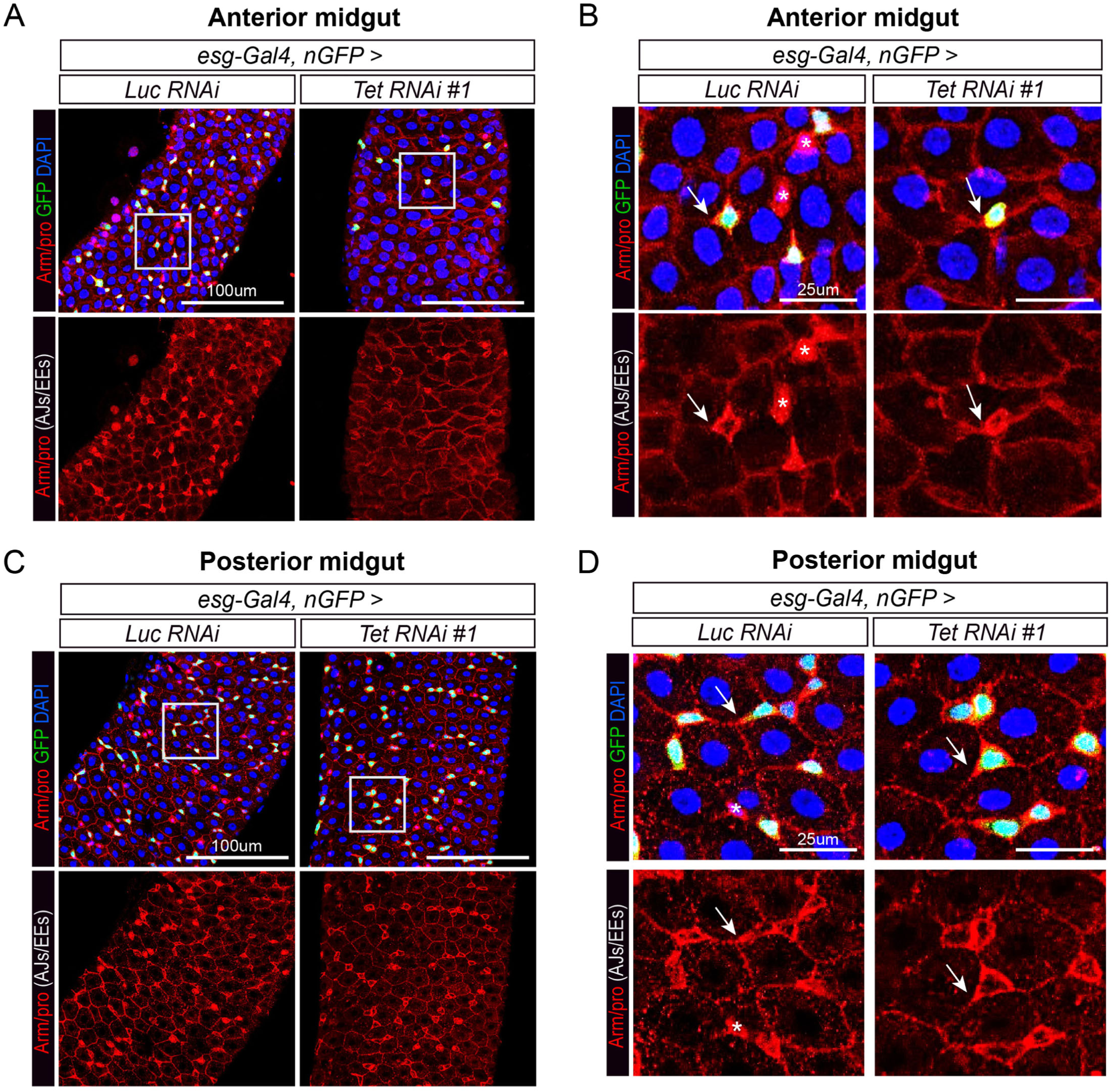
Developmental loss of *Tet* causes structural defects in the adult midgut. (A) Representative images of anterior midguts from control (*Luc RNAi*) and *Tet RNAi* #1 adults, demonstrating structural defects resulting from *Tet* depletion. (B) High-magnification views of the insets in (A). Arrows indicate tricellular junctions, and asterisks denote enteroendocrine (EE) cells. Control epithelia exhibit sharply defined cell boundaries, whereas *Tet RNAi* midguts display severely disorganized and poorly defined edges. (C) Representative images of posterior midguts from the same genotypes, showing no obvious structural or junctional defects. (D) High-magnification views of the insets in (C), with arrows marking normal, intact tricellular junctions.

**Figure S3.**
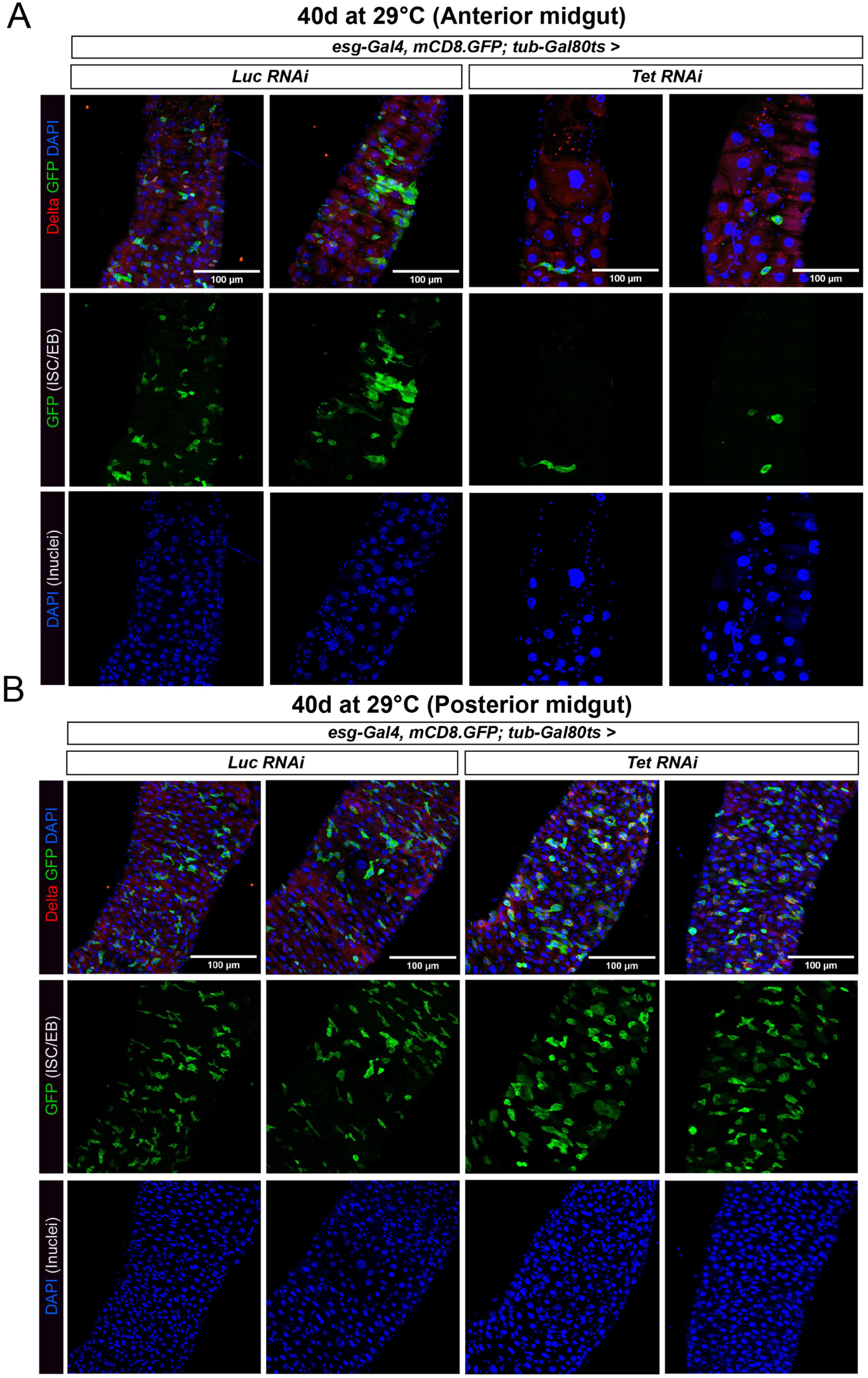
Adult-specific loss of *Tet* induces anterior progenitor loss and EC compensatory hypertrophy. (A) Representative images of anterior midguts from control (*Luc RNAi*) and *Tet RNA*i flies 40 days after adult-specific induction. *Tet*-depleted adults exhibit a severe depletion of the ISC/EB pool (green) accompanied by compensatory hypertrophy of the surrounding ECs. (B) Representative images of posterior midguts from the same cohort 40 days post-induction, demonstrating no obvious morphological differences or progenitor loss between controls and *Tet RNAi*.

**Figure S4.**
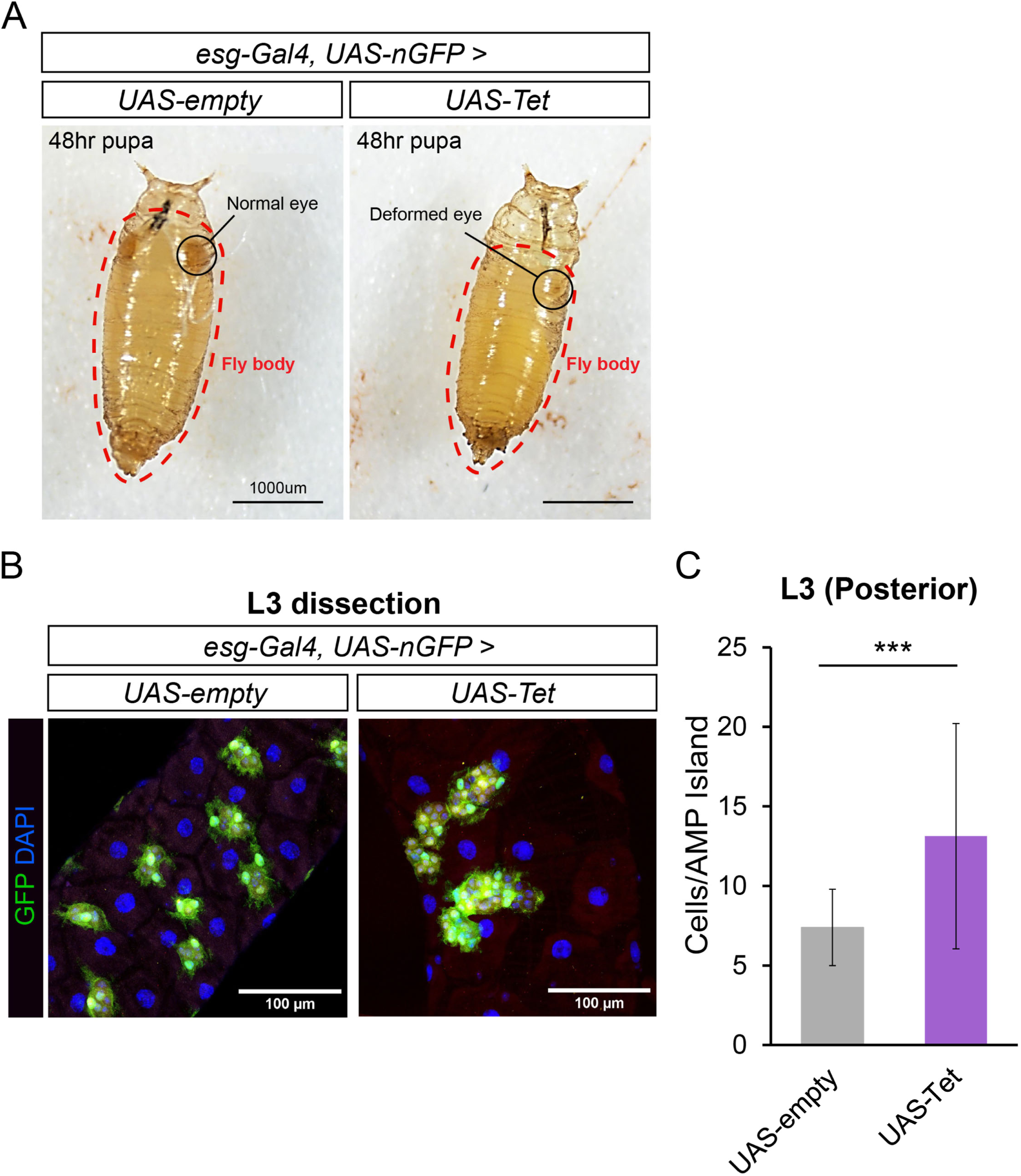
Developmental overexpression of *Tet* induces mid-pupal lethality and hyperproliferation of larval AMPs. (A) Representative images of 48hr pupae within their puparia. Control pupae (left) exhibit normal development, whereas *Tet* overexpression (right) results in mid-pupal lethality, evidenced by a lack of eye development and widespread tissue degeneration. (B) Representative confocal images of posterior midguts from L3 control (*UAS-empty*) and *Tet* overexpression (*Tet* OE) larvae. *Tet* OE drives a massive expansion of the progenitor pool, resulting in a significant increase in the number of cells per AMP island, consistent with the hyperproliferation observed in the anterior midgut. (C) Quantification of the number of cells per AMP island in control versus *Tet* OE midguts. Bars represent mean ± SD. \**p* ≤ 0.05, \*\**p* ≤ 0.01, \*\*\**p* ≤ 0.001. *p* values were from unpaired Student’s t-test.

**Figure S5.**
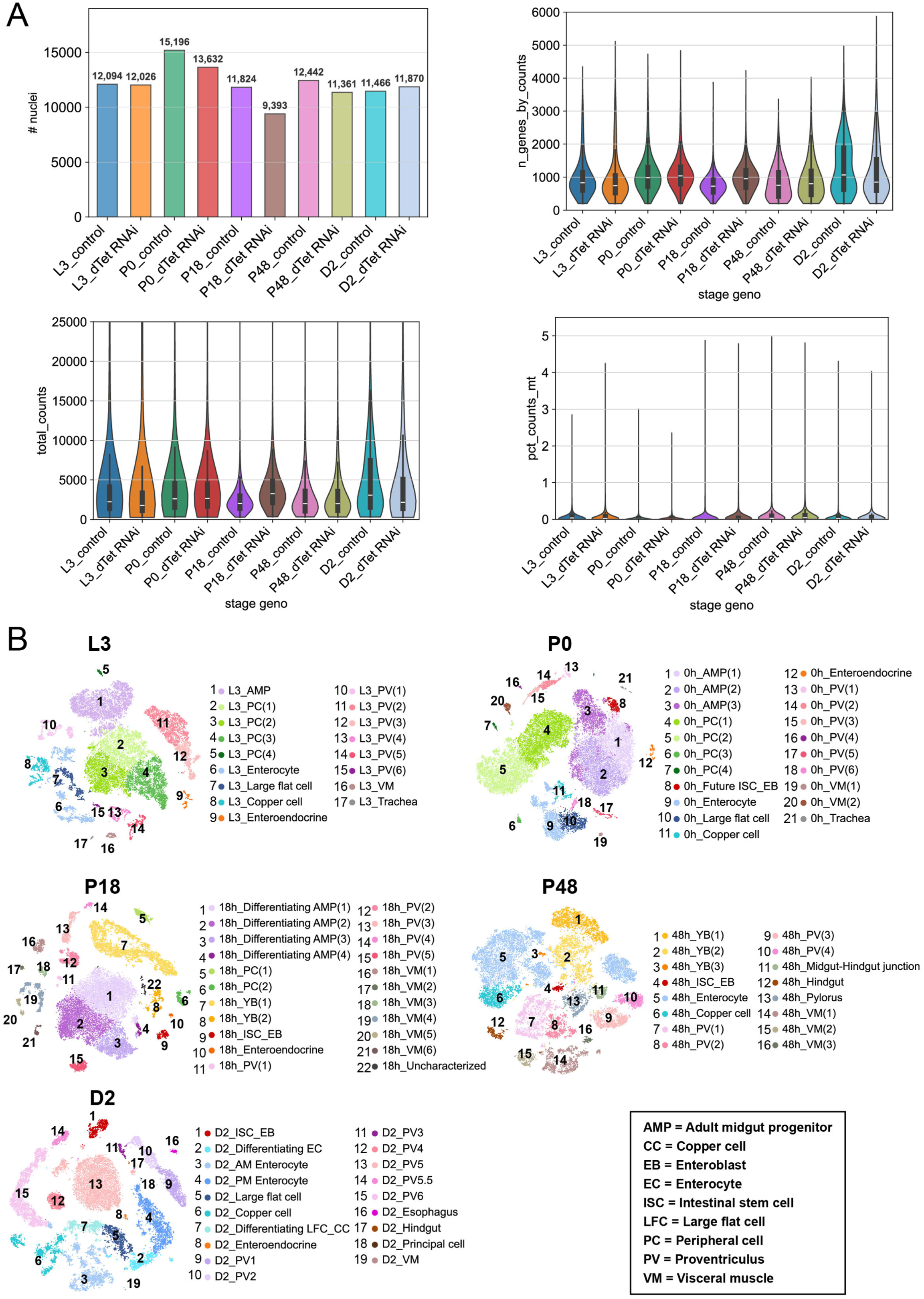
**snRNA-seq quality control metrics and detailed stage-specific annotations.** (A) Number of nuclei, expressed gene count, UMI counts, and mitochondria transcript ratios of each sample for the gut atlas dataset. (B) Detailed annotation across the five stages of midgut development.

**Figure S6.**
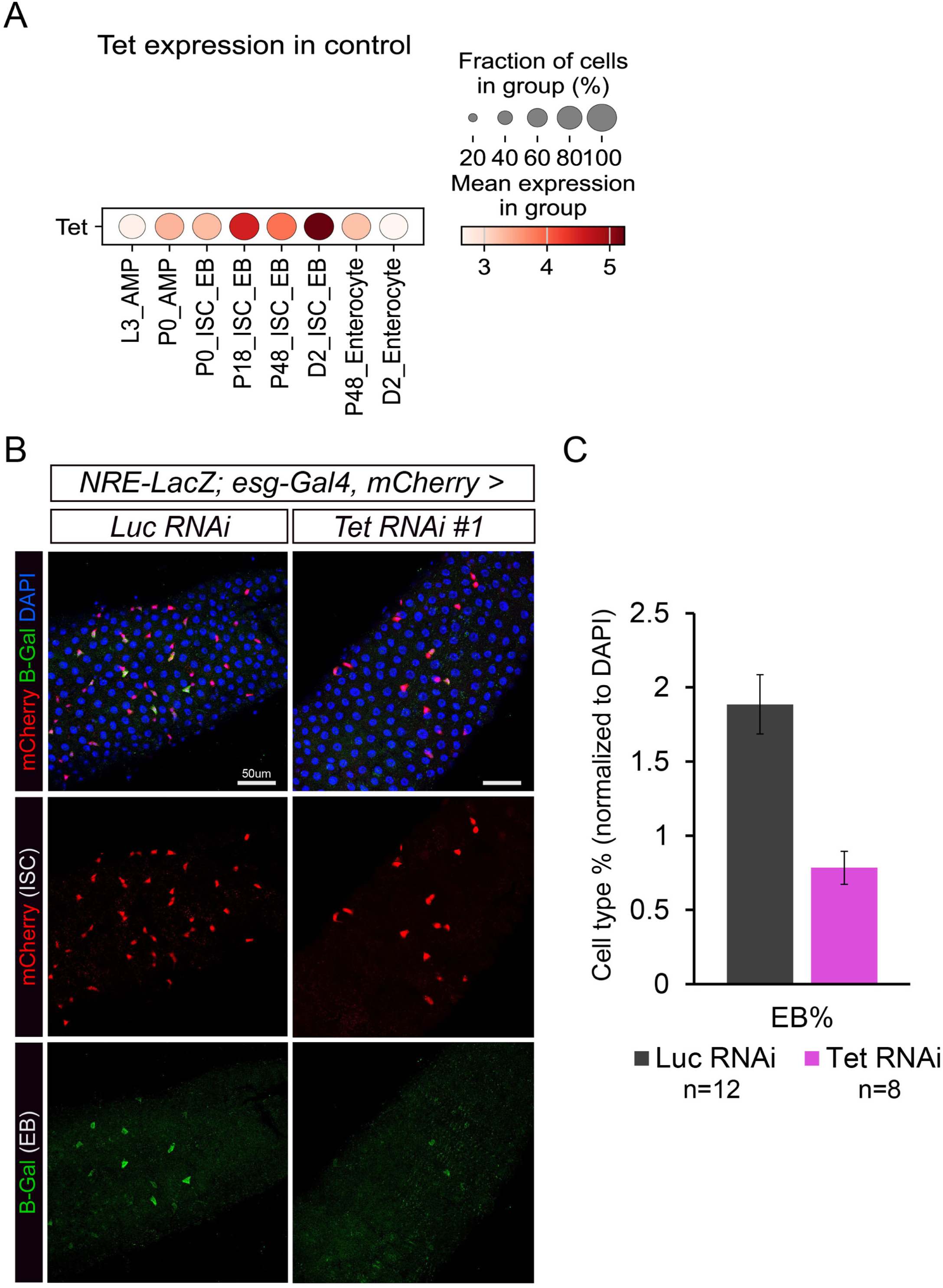
Loss of *Tet* promotes rapid transition through the EB phase. (A) Representative images of young adult (day 1) midguts from control and *Tet RNAi* flies, distinguishing ISCs (red) from transit-amplifying EBs (green). (B) Quantification of NRE-lacZ+ EBs. The lack of accumulating EBs combined with the loss of ISCs (Fig. 1D) suggests rapid transition through the EB stage. This supports a model in which *Tet*-depleted progenitors fail to maintain a stable intermediate state and instead undergo rapid, premature differentiation.

**Table S1.** Identification of stage-specific midgut lineage markers and their temporal dysregulation following *Tet* depletion. This table compiles the broad and stage-specific marker genes used to annotate the developmental trajectories of Adult Midgut Progenitors (AMPs), Intestinal Stem Cells/Enteroblasts (ISC/EBs), and Enterocytes (ECs). For each control cluster, the top 30 defining differentially expressed genes (DEGs) are provided. To map the progressive transcriptional collapse induced by *Tet* loss, the differential expression (Log2 Fold Change) between *Tet RNAi* and control nuclei is overlaid for each marker gene across the captured developmental stages (L3 to Day 2 Adult). Blank cells within the *Tet RNAi* columns denote comparisons that did not reach statistical significance (adjusted *p* ≥ 0.05).

**Table S2.** *In vivo* functional screen of candidate *Tet* transcriptional targets. 51 fly lines for 25 candidate genes were selected based on significant differential expression in *Tet RNAi* ISC/EB populations at the P18 developmental stage compared to controls. Transcriptomic statistics (Log2 Fold Change and Adjusted p-value) are derived from snRNA-seq analysis of the P18 timepoint. To determine if dysregulation of a single downstream target was sufficient to phenocopy the *Tet RNAi*, indicated UAS-RNAi or UAS-overexpression (O/E) lines were crossed to the progenitor-specific *esg-Gal4, UAS-nGFP* driver at 25°C. Adult midguts were dissected and qualitatively scored via epifluorescence microscopy specifically for the presence or absence of a profound, penetrant loss of the GFP-positive progenitor pool. "Lethal / Unscorable" designates crosses that result in death before eclosion.

